# SARS-CoV-2 vaccination induces immunological memory able to cross-recognize variants from Alpha to Omicron

**DOI:** 10.1101/2021.12.28.474333

**Authors:** Alison Tarke, Camila H. Coelho, Zeli Zhang, Jennifer M. Dan, Esther Dawen Yu, Nils Methot, Nathaniel I. Bloom, Benjamin Goodwin, Elizabeth Phillips, Simon Mallal, John Sidney, Gilberto Filaci, Daniela Weiskopf, Ricardo da Silva Antunes, Shane Crotty, Alba Grifoni, Alessandro Sette

## Abstract

We address whether T cell responses induced by different vaccine platforms (mRNA-1273, BNT162b2, Ad26.COV2.S, NVX-CoV2373) cross-recognize SARS-CoV-2 variants. Preservation of at least 83% and 85% for CD4^+^ and CD8^+^ T cell responses was found, respectively, regardless of vaccine platform or variants analyzed. By contrast, highly significant decreases were observed for memory B cell and neutralizing antibody recognition of variants. Bioinformatic analyses showed full conservation of 91% and 94% of class II and class I spike epitopes. For Omicron, 72% of class II and 86% of class I epitopes were fully conserved, and 84% and 85% of CD4^+^ and CD8^+^ T cell responses were preserved. In-depth epitope repertoire analysis showed a median of 11 and 10 spike epitopes recognized by CD4^+^ and CD8^+^ T cells from vaccinees. Functional preservation of the majority of the T cell responses may play an important role as a second-level defense against diverse variants.

## INTRODUCTION

The emergence of numerous SARS-CoV-2 variants of interest (VOI) and of concern (VOC) is one of the most important developments in the COVID-19 pandemic (Callaway, 2021). Our understanding of the virological and immunological features associated with the main VOCs is key to inform health policies, including boosting and vaccination schedules, and also inform the development of potential variant-specific or pan-coronavirus vaccines. Important aspects include whether the different variants are more infectious, more easily transmissible, linked to more severe disease, and escape immune responses induced by either vaccination or natural infection.

The Alpha (B.1.1.7), Beta (B.1.351) and Gamma (P.1) VOCs were reported in the late 2020-May 2021 period (Harvey et al., 2021; Walensky et al., 2021). Several additional variants were described more recently (May-Oct 2021)(Chakraborty et al., 2021; Otto et al., 2021), including Mu (B.1.621)(Uriu et al., 2021) and Delta (B.1.617.2)(Mlcochova et al., 2021), with the latter quickly becoming the most dominant SARS-CoV-2 lineage worldwide. Omicron (B.1.1.529) is the latest VOC, reported in November 2021, and stands out for the larger number of spike mutations compared to other VOCs (Karim and Karim, 2021), its transmissibility even in the presence of Delta, and its ability to spread in populations with high levels of immunity. It is expected to become dominant globally in the coming weeks.

A number of knowledge gaps remain in terms of our understanding of VOI/VOCs in relation to T and B cell immune reactivity. While the impact of the variant-associated mutations has been established for most variants in terms of antibody reactivity (Garcia-Beltran et al., 2021; Stamatatos et al., 2021), including studies on Omicron (Liu et al., 2021; Planas et al., 2021; Schmidt et al., 2021), much less is available for memory T cells and memory B cells. Memory T cell and B cell recognition of variants are important issues. Several lines of evidence point to potential roles of T cells in modulating disease severity and contributing to disease protection (Gagne et al., 2021; Sette and Crotty, 2021; Tan et al., 2021). The continued maturation of B cell responses over time (Cho et al., 2021; Dan et al., 2021; Goel et al., 2021) may play an important role in adapting SARS-CoV-2 immunity to VOCs. Regarding memory T cells, we and others previously demonstrated that for the early variants Alpha, Beta, Gamma and Epsilon the impact of mutations is limited and the majority of CD4^+^ and CD8^+^ T cell responses are preserved in both vaccinated and natural infection responses (Collier et al., 2021; Geers et al., 2021; Keeton et al., 2021; Melo-Gonzalez et al., 2021; Riou et al., 2021; Tarke et al., 2021b).

However, studies on the impact of newer variants on T cells, including Mu and Omicron in particular are limited or missing (Madelon et al., 2021). If the majority of T cell responses are maintained, the T cells may play an important role as a second line of defense, in light of the substantial escape from antibody responses. In this study, we focus on a large panel of variants to understand the impact of more recent variants on T cell responses and B cell memory as compared to the early variants, particularly in the context of COVID-19 vaccination and evaluation of the adaptive responses induced by different vaccine platforms.

## RESULTS

### Cohort of COVID-19 vaccinees to assess T cell responses to a panel of SARS-CoV-2 variants

To assess the cross-recognition capability of T cell responses induced by different vaccine platforms, we enrolled a cohort of 96 adult individuals vaccinated with different vaccines currently in use in the United States under FDA EUA: mRNA-based mRNA-1273, mRNA based BNT162b2 and the adenoviral vector-based Ad26.COV2.S. Subjects immunized with a recombinant protein recombinant vaccine NVX-CoV2373, currently approved in the EU and in clinical assessment for the US, were also studied. To determine the longevity of T cell cross-recognition of the different SARS-CoV-2 variants, we studied samples from four different time points; 2 weeks after 1^st^ dose of vaccination, 2 weeks after 2^nd^ dose of vaccination, 3.5 months, and 5-6 months after the last vaccination dose received. Based on sample availability, the study design was cross-sectional. One control donor cohort was also enrolled of early convalescent donors who had mild disease (collected approximately one month post symptom onset, range 21-43 days).

Characteristics of the donor cohorts analyzed are summarized in **Table S1**. The sub-cohorts were approximately matched for gender and age across time points. For each time point, the days post-vaccination (dPV) and the SARS-CoV-2 S protein RBD immunoglobulin G (IgG) ELISA titers are detailed as a function of the vaccine platform analyzed and the time point of sample collection. In addition, nucleocapsid (N) IgG was also run to assess previous infection, with the highest frequency of positive response of 14% observed at time point 3 in Ad26.COV2.S recipients (**Table S1**). HLA typing for the vaccinated cohort is presented in **Table S2**.

We previously reported T cell reactivity to Alpha, Beta and Gamma variants of concern (Tarke et al., 2021b). Since then, by July 2021 an additional 5 SARS-CoV-2 VOI/VOCs emerged, namely B.1.1.519, Kappa (B.1.617.1), Delta (B.1.617.2), Lambda (C.37) and R.1. To estimate the impact of the different variants on T cell responses after vaccination, we mapped the specific spike protein mutations (amino acid replacements and deletions) as compared to the SARS-CoV-2 Wuhan ancestral sequence (**Table S3**). For the Wuhan ancestral sequence and each of the variants analyzed, we generated MegaPools (MP) of 15-mer peptides, overlapping by 10 amino acids, spanning the entire spike protein.

### Spike-specific CD4^+^ and CD8^+^ T cell reactivity to different SARS-CoV-2 variants in fully vaccinated individuals

Next, we evaluated T cells from the vaccine cohorts for their capacity to cross-recognize MPs spanning the entire spike sequences of different variants, compared to a control MP spanning the ancestral spike antigen. First, T cell responses were determined from blood samples of fully immunized subjects two weeks after the second immunization to the mRNA-based vaccines mRNA-1273 and BNT162b2, and 6 weeks post-immunization with the adenoviral vector-based Ad26.COV2.S. To measure the T cell responses, we combined activation-induced marker (AIM) assays (Tarke et al., 2021b) with cytokine intracellular staining (ICS) (Mateus et al., 2021). A comparison of the AIM and ICS protocols performed separately with the AIM+ICS combined protocol showed no significant differences in the markers analyzed (**Figure S1A-C**).

CD4^+^ and CD8^+^ T cell responses to spike MPs derived from the ancestral strain and from the corresponding MPs representing Alpha, Beta, Gamma, Delta, B.1.1.519, Kappa, Lambda and R.1 variants, were measured by AIM OX40^+^CD137^+^ and CD69^+^CD137^+^ for CD4^+^ and CD8^+^ T cells, respectively (Tarke et al., 2021b). For each subject/variant/vaccine combination, we calculated the T cell response fold change relative to the ancestral sequence (variant/ancestral). Only donors with a positive spike ancestral response were included in the analysis (CD4: LOS= 0.03%, SI>2; CD8: LOS= 0.04%, SI>2). **Figure 1** summarizes the fold change results for all vaccine platforms combined and separately, for CD4^+^ (**Figure 1B**) and CD8^+^ (**Figure 1D**) T cell responses. For all variants, regardless of the vaccine platform considered, no significant decrease (fold change < 1.00 by the Wilcoxon Signed Rank T test compared to a hypothetical median of 1) was detected. In all cases, the geomean fold change variation was close to 1.00 (i.e., no change). The average fold change values considering all 24 different vaccine/variant combinations (3 vaccine platforms and 8 variants) was 1.01 (range 0.84 to 1.3) for CD4^+^ and 1.1 (range 0.81 to 1.5) for CD8^+^ T cells. At the level of individual donors, a decrease greater than an arbitrary 3-fold threshold (0.3 fold-change, indicated by dotted lines) was only observed for two Ad26.COV2.S vaccinees and the Lambda variant, one for CD4^+^ and another donor for CD8^+^ T cells. No variant/donor combination was associated with decreases greater than 10-fold. **Figure S2** shows the corresponding AIM^+^ percentages and their relative paired comparisons based on the magnitude of the responses for each variant with the ancestral spike reactivity. Analysis of the Coefficient of Variation (CVs) of the fold changes for each variant across vaccine platforms revealed significant differences in the variation across vaccine platforms (CD4^+^: mRNA-1273 Vs Ad26.COV2.S P=0.0009; BNT162b2 Vs Ad26.COV2.S P= 0.0078; CD8^+^ : mRNA-1273 Vs Ad26.COV2.S P=0.0024; BNT162b2 Vs Ad26.COV2.S P=0.1230; Mann-Whitney with multiple comparison correction). Overall, these results indicate that both CD4^+^ and CD8^+^ T cell responses are largely conserved, irrespective of the variant and vaccine platform considered.

**Figure 1.**
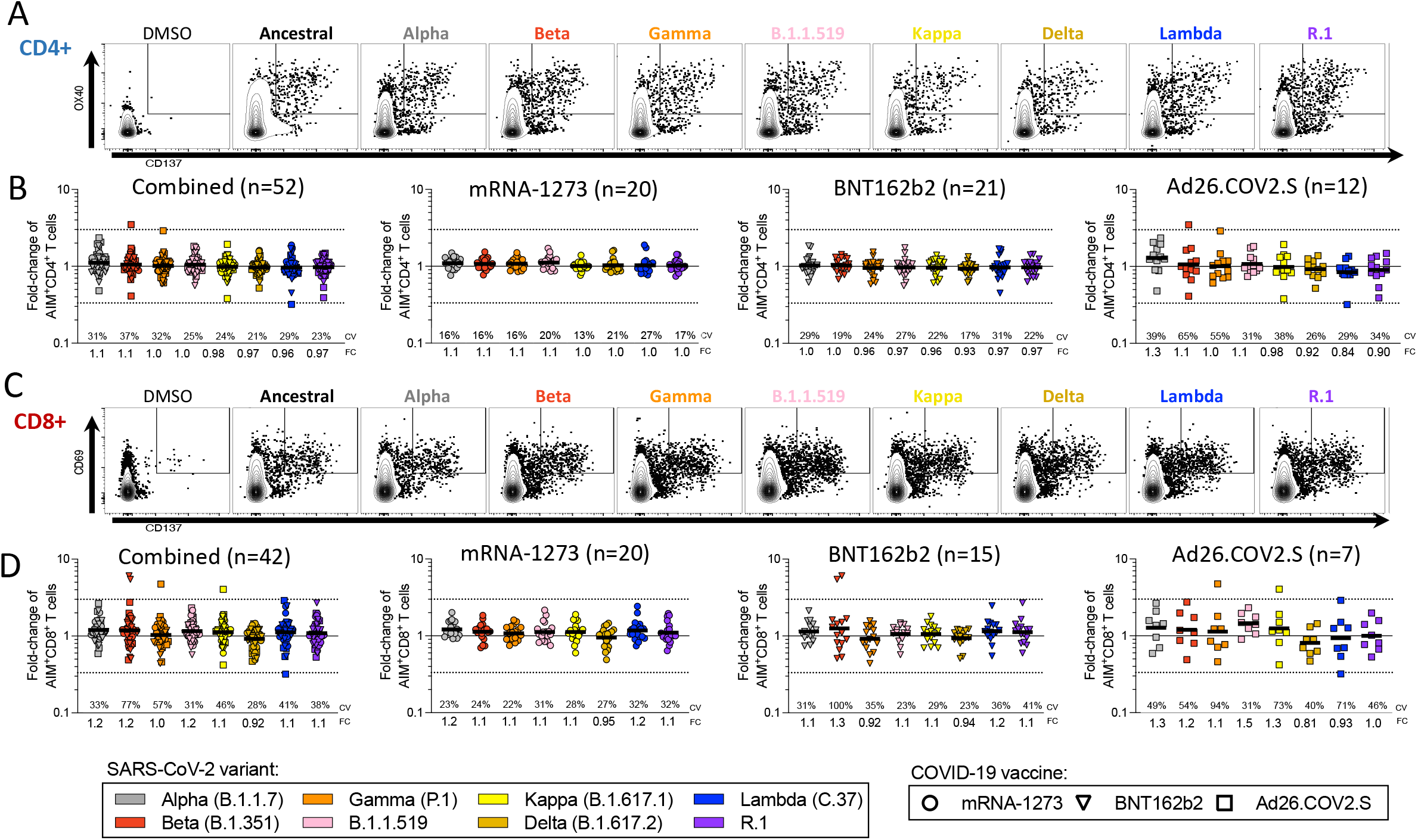
Impact of variant associated mutations on spike-specific CD4^+^ and CD8^+^ T cell responses to SARS-CoV-2 variants. T cell responses of PBMCs from fully vaccinated COVID-19 vaccinees were assessed with variant spike MPs. The effect of mutations associated with each variant MP is expressed as relative (fold change variation) to the T cell reactivity detected with the ancestral strain MP. Results from COVID-19 mRNA-1273 (circles), BNT162b2 (triangles) and Ad26.COV2.S (squares) vaccinees are presented combined together, and separately by vaccine platform. For fold change calculations, only donors responding to the ancestral S MP were considered. (**A**) Representative gating of CD4^+^ T cells of a mRNA-1273 vaccine recipient responding to different SARS-CoV-2 variants MPs. (**B**) Fold-change is calculated for AIM^+^ CD4^+^ T cells relative to the ancestral strain in COVID-19 vaccinees. (**C**) Representative gating example of a mRNA-1273 vaccine recipient for CD8^+^ T cells against the SARS-CoV-2 variants in analysis. (**D**) Fold change is calculated for AIM^+^ CD8^+^ T cells relative to the ancestral strain in COVID-19 vaccinees. Coefficients of variation (CV) for the variants are listed in each graph. Significance of fold change decreases for each variant was assessed by Wilcoxon Signed Rank T test compared to a hypothetical median of 1. See also Figures S1, S2 and S3 and Table S1.

By ICS, significant CD4^+^ T cell responses to the ancestral Wuhan spike pool were observed for 48 subjects and CD8^+^ T cell responses were observed for 24 subjects. Thus, combined ICS results for all vaccine platforms are presented. CD4^+^ T responses were associated with a polyfunctional response, encompassing IFNγ, TNFα, IL-2, granzyme B, or CD40L (**Figure 2A-B, Figure S1**). Cytokine^+^ CD8^+^ T cells were measured using IFNγ, TNFα, IL-2, or granzyme B (**Figure 2C-D, Figure S1**). Overall, the average fold change values considering all individual variant combinations was 1.07 (range 0.96 to 1.1) for CD4^+^ and 0.97 (range 0.75 to 1.3) for CD8^+^ T cells, with the greatest decrease of a fold change of 0.75 geomean seen for Delta for cytokine^+^ CD8^+^ T cells. At the level of individual CD4^+^ T cell responses, a decrease greater than 3-fold was observed for one donor with the Beta variant (**Figure 2A**) and no decreases greater than 10-fold were observed. For CD8^+^ T cells, decreases greater than 10-fold were observed for one Ad26.COV2.S donor with the Alpha, B.1.1.519, and R.1 variants and one mRNA-1273 donor with Delta. Decreases in the 3-10 fold range were observed for one BNT162b2 donor with Gamma and Lambda, and for two different donors for BNT162b2 with Kappa, and Ad26.COV2.S with Lambda (**Figure 2C**).

**Figure 2.**
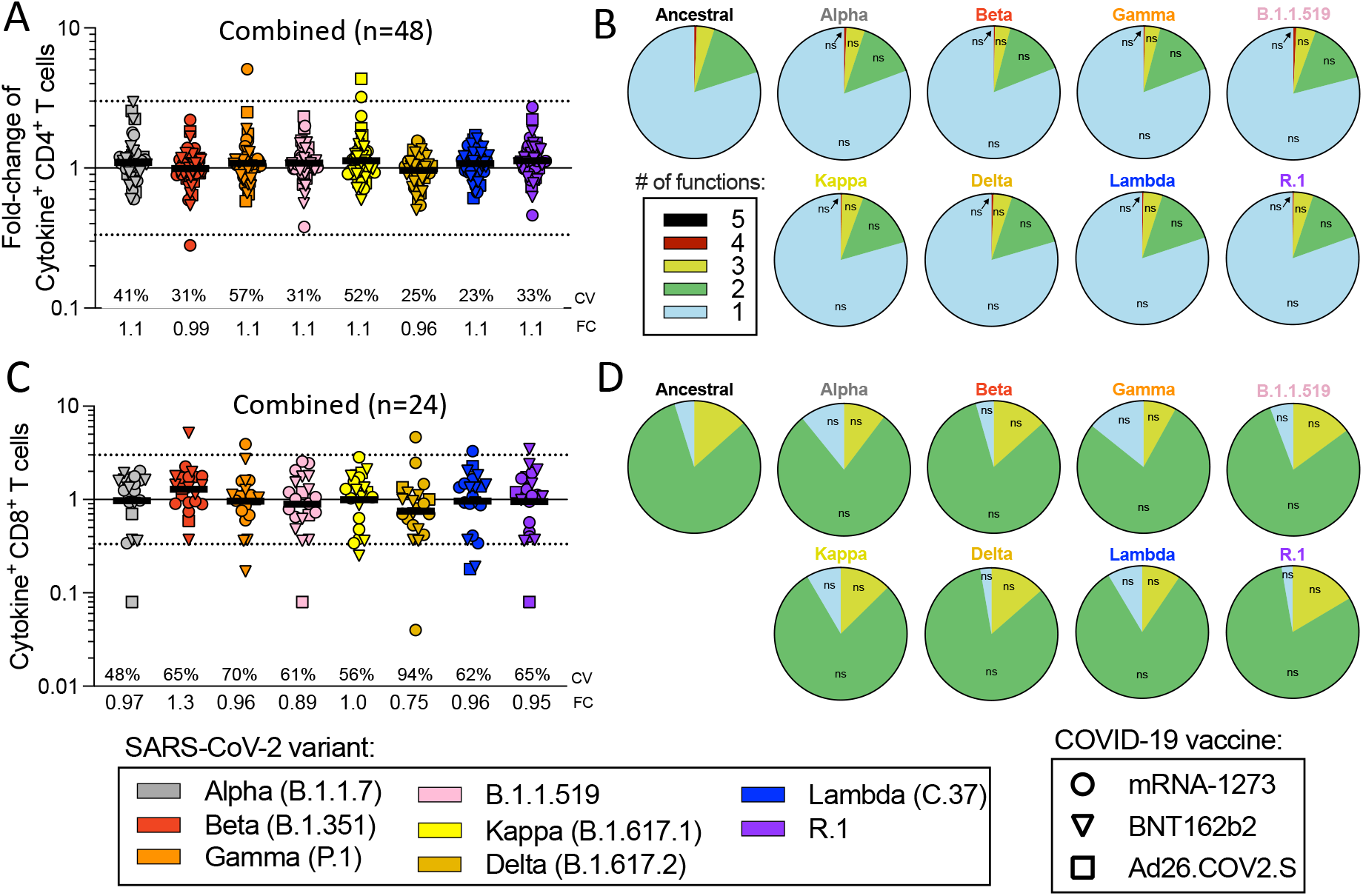
Impact of variant associated mutations on spike-specific cytokine responses in CD4^+^ and CD8^+^ T cells. Fully vaccinated COVID-19 vaccinees were assessed with variant spike MPs and the effect of mutations associated with each variant MP is expressed as relative (fold change variation) to the T cell reactivity detected with the ancestral strain MP. Results from COVID-19 mRNA-1273 (circles), BNT162b2 (triangles) and Ad26.COV2.S (squares) vaccinees are presented combined together. (**A**) Fold change values for cytokine^+^CD4^+^ T cells are calculated based on the sum of CD4^+^ T cells producing CD40L, IFNγ, TNFα, IL-2, or Granzyme B and (**B**) the functionality of the CD4^+^ T cell is calculated by looking at the different combinations of cytokines. (**C**) Fold change values for cytokine^+^CD8^+^ T cells are calculated based on the sum of CD8^+^ T cells producing IFNγ, TNFα, IL-2, or Granzyme B and (**D**) the functionality of the CD8^+^ T cell is calculated by looking at the different combinations of cytokines. Coefficients of variation (CV) for the variants are listed in each graph. Significance of fold change decreases for each variant was assessed by Wilcoxon Signed Rank T test compared to a hypothetical median of 1. See also Figures S1 and S2 and Table S1.

Considering the different assay readouts (AIM and ICS) and different donors analyzed, the fold change was calculated in 166 instances for 8 different variants, for a total of 1328 determinations. T cell responses with decreases greater than 3-fold were observed in 11 instances (0.82%) of variant/subject combinations tested, and decreases greater than 10-fold were observed in 4 instances (0.30%) of variant/subjects tested. Thus, in almost 99% of cases the differences were less than 3-fold.

When we measured AIM^+^ T cells responses at time point 1 (**Figure S3**), after a single dose of vaccine, we found a very similar pattern to what was observed in time point 2, with no significant decreases in each of the variants analyzed at both population and individual level, except for 3 out of 280 instances (1%) with a 3-fold decrease in CD8^+^ AIM+ T cells (**Figure S3B**). The overall geomean fold change reactivity, averaging all donors from all vaccine platforms in time point 2, for the Delta variant was 0.97 for CD4^+^ T cells by AIM and 0.96 by ICS. For CD8^+^ T cells, the overall geomean fold change was 0.82 by AIM and 0.80 by ICS. Cytokine-based readouts might be relatively more impacted by variant mutations. These results confirm, in a larger dataset, that T cell responses from vaccinated subjects are largely preserved against Alpha, Beta and Gamma (Tarke et al., 2021b). Importantly, these results extend these observations to more VOI/VOCs, including the prominent Delta variant.

### Cross-recognition of SARS-CoV-2 variants by memory T cell responses

We then examined memory T cells and memory B cells 3-4 months after vaccination. At this time point, samples from a number of NVX-CoV2373 vaccinated individuals were available and therefore included in the analysis. Spike-specific CD4^+^ T cell memory was characterized by AIM and ICS (**Figure 3A-C, Figure S4A-B**), including memory circulating T follicular helper cells (cT_FH_) (**Figure 3G, Figure S4E**).

**Figure 3.**
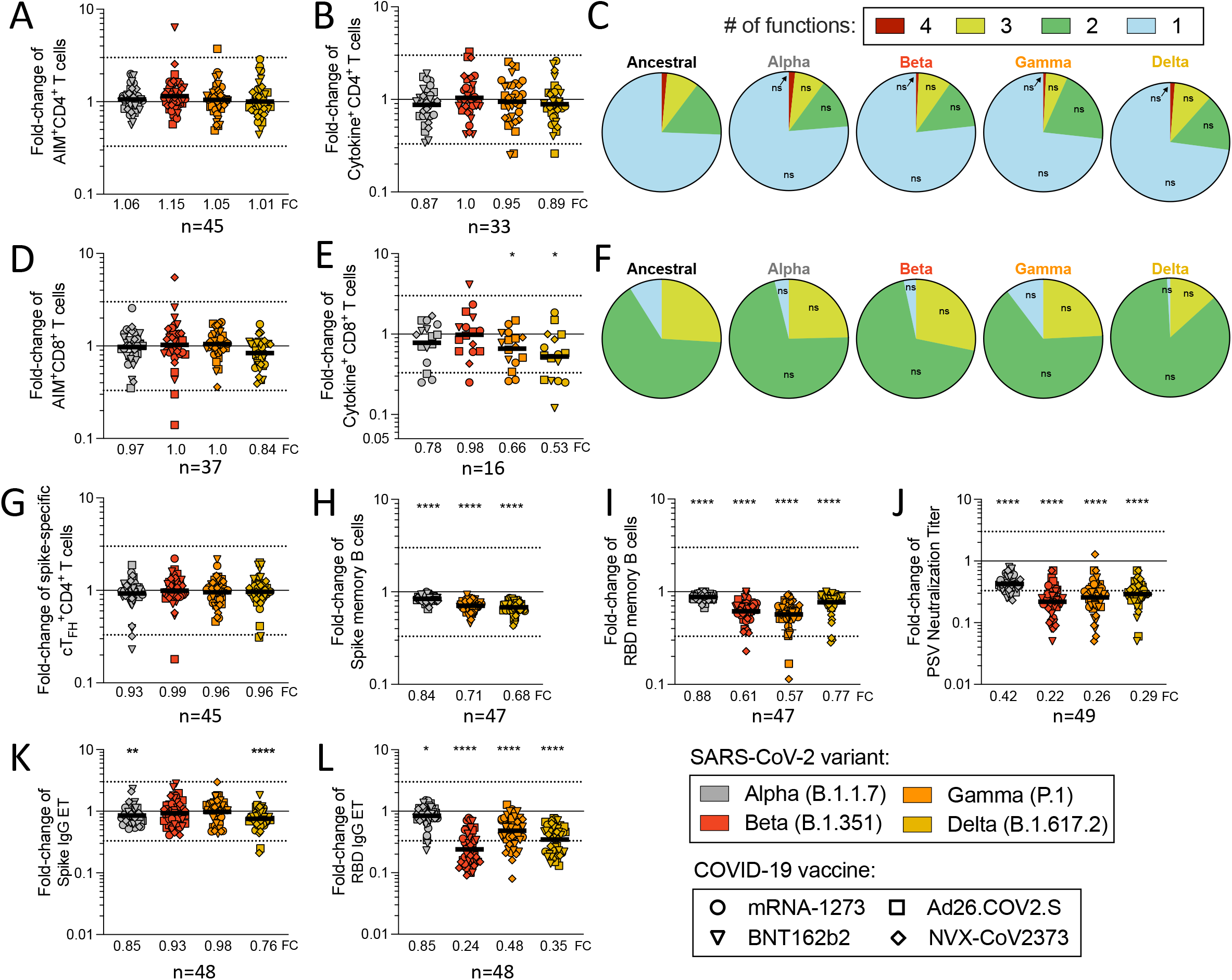
Vaccinee memory T and B cell recognition of COVID-19 variants. Fully vaccinated recipients of the COVID-19 mRNA-1273 (circles), BNT162b2 (triangles), Ad26.COV2.S (squares) and NVX-CoV2373 (diamonds) vaccines were assessed for T and B cell responses to variant Spikes. Fold-change values were calculated based on the response to the ancestral S when there was a measurable response. Fold change values are shown for CD4^+^ by (**A**) AIM and (**B**) ICS assay. (**C**) The functional profile of cytokine producing CD4^+^ T cells was calculated as the percentage of cells with 1, 2, 3, or 4 functions defined by intracellular staining for IFNγ, TNFα, IL-2, Granzyme B, or CD40L. For CD8^+^ T cells, the fold change values are shown for the (**D**) AIM and (**E**) ICS assay. (**F**) The functional profile of cytokine producing CD8^+^ T cells was calculated as the percentage of cells with 1, 2, 3, or 4 functions defined by intracellular staining for IFNγ, TNFα, IL-2, or Granzyme B. p values for the functional profile of CD4^+^ and CD8^+^ T cells were calculated by Mann-Whitney. (**G**) Spike-specific cT_FH_^+^CD4^+^ T cells were calculated based on the CXCR5 expression of the AIM^+^CD4^+^ T cells. Memory B cell responses are shown corresponding to (**H**) spike and (**I**) RBD. Fold change values are shown for the (**J**) antibody neutralization assay as well as (**K**) spike and (**L**) RBD IgG serology. The geometric mean is listed at the bottom of the memory B cell and serology graphs. Significance of fold change decreases for each variant was assessed by Wilcoxon Signed Rank T test compared to a hypothetical median of 1. See also Figures S1 and S4 and Table S1.

No significant decrease of memory CD4^+^ T cell recognition of Alpha, Beta or Gamma variants was observed by AIM, cytokine or cT_FH_ metrics (**Figure 3A-B, G**). Mean preservation of CD4 reactivity was 0.91 (range 0.87 to 1.2) considering all three assays and variants. At the individual level, no substantial decreases in CD4^+^ T cell variant recognition were observed by AIM. Cytokine response decreases >3-fold were observed in 3 out of 144 instances (2%). cT_FH_ memory cell recognition of variants decrease >3-fold in 5 (3.5%) instances (**Figure 3G**).

Spike-specific CD8^+^ T cell memory was characterized by overall geomean fold change of 0.95 by AIM and 0.74 by ICS (**Figure 3D-F, Figure S4C-D**). No significant decrease of memory CD8^+^ T cell recognition of Alpha, Beta or Gamma variants was observed by AIM. Decreases of 0.66 and 0.53 fold change were observed for Gamma and Delta, respectively, for memory CD8^+^ T cell by ICS (**Figure 3D-F**). At the individual level, decreases >3-fold were observed in 2 out of 148 instances (1.3%) of variant recognition by AIM. Cytokine response decreases >3-fold were observed in 12 out of 64 instances (18.8%), none of which were greater than 10-fold (2%). Thus, the overall pattern of variant recognition for memory CD4^+^ and CD8^+^ T cells paralleled the peak T cell responses (**Figures 1–3**). Memory CD4^+^ T cell recognition of these variants was largely preserved, including Delta. Some memory CD8^+^ T cell recognition decreases were noted by cytokine production, most notably against Delta for which a 1.9-fold decrease was found.

We next examined the ability of spike-specific memory B cells to recognize variants Alpha, Gamma and Delta (**Figure 3H, Figure S4F,H**). Significant losses in memory B cell recognition of spike for Alpha (variant/Wuhan fold change=0.84; P<0.0001), Gamma (fold change=0.71; P<0.0001) and Delta (fold change=0.68; P<0.0001) were observed (**Figure 3H**). The receptor binding domain (RBD) of spike is the primary target of SARS-CoV-2 neutralizing antibodies and a site of variant neutralizing antibody escape. We therefore characterized RBD-specific memory B cell recognition of variants Alpha, Beta, Gamma and Delta (**Figure 3I, Figure S4G,I**). Significant decreases in RBD-specific memory B cell recognition of Alpha (fold change=0.88; P<0.0001), Beta (fold change=0.61; P<0.0001), Gamma (fold change=0.57; P<0.0001) and Delta (fold change=0.77; P<0.0001) were all noted (**Figure 3I**).

Neutralizing antibody titers to variants were measured compared to a D614G reference virus, for the same vaccinated individuals (**Figure 3J, Figure S4J**). Neutralization decreases were significant for Alpha (P<0.0001), Beta (P<0.0001), Gamma (P<0.0001) and Delta (P<0.0001) variants (**Figure 3J**). The highest neutralization antibody titers were against D614G, and reductions in neutralizing titers of 2.4-fold, 4.5-fold, 3.8-fold and 3.4-fold against Alpha, Beta, Gamma and Delta variants, respectively were noted (**Figure 3J** and **Figure S4J**). A similar pattern was observed for COVID-19 convalescent subjects (**Figure S4K-L**). Spike and RBD binding IgG titers in vaccinated subjects had similar trends to neutralizing antibodies but with smaller differences (**Figure 3K-L, Figure S4M-N**). In conclusion, while no significant change in T cell recognition was noted, decreases in memory B cell and neutralizing antibody recognition of all variants analyzed were apparent.

### Predicted impact of SARS-CoV-2 variants on T cell epitopes

With the recent emergence of the Omicron variant, studies were immediately expanded to include Omicron. We first predicted the impact of variant mutations for CD4^+^ and CD8^+^ T cell epitopes experimentally curated in the IEDB (www.IEDB.org)(Grifoni et al., 2021; Vita et al., 2019) (**Table S4**). In addition to Omicron, we included a wider panel of early and late SARS-CoV-2 variants for comparison.

For CD4^+^ T cell responses, an average of 95% of the epitopes spanning the entire SARS-CoV-2 proteome were fully conserved (no mutations) across the variants (**Figure 4A**). The Delta variant was not associated with a significant decrease (**Figure 4A**), while the fraction of fully conserved epitopes was reduced in Omicron (88%), compared to the other variants (p<0.0001) (**Figure 4A**). A similar result was observed for CD8^+^ T cell epitopes, with an average 98% overall conservation but 95% for Omicron (P<0.0001) (**Figure 4B**). Considering only spike epitopes, of relevance in the context of vaccination, an average of 91% and 94% CD4^+^ and CD8^+^ T cell epitopes were conserved in the various variants. The Omicron variant was associated with the fewest fully conserved spike epitopes for both CD4^+^ and CD8^+^ T cells (CD4^+^: 72%, P<0.0001; CD8^+^: 86%, P<0.0001) (**Figure 4C-D**).

**Figure 4.**
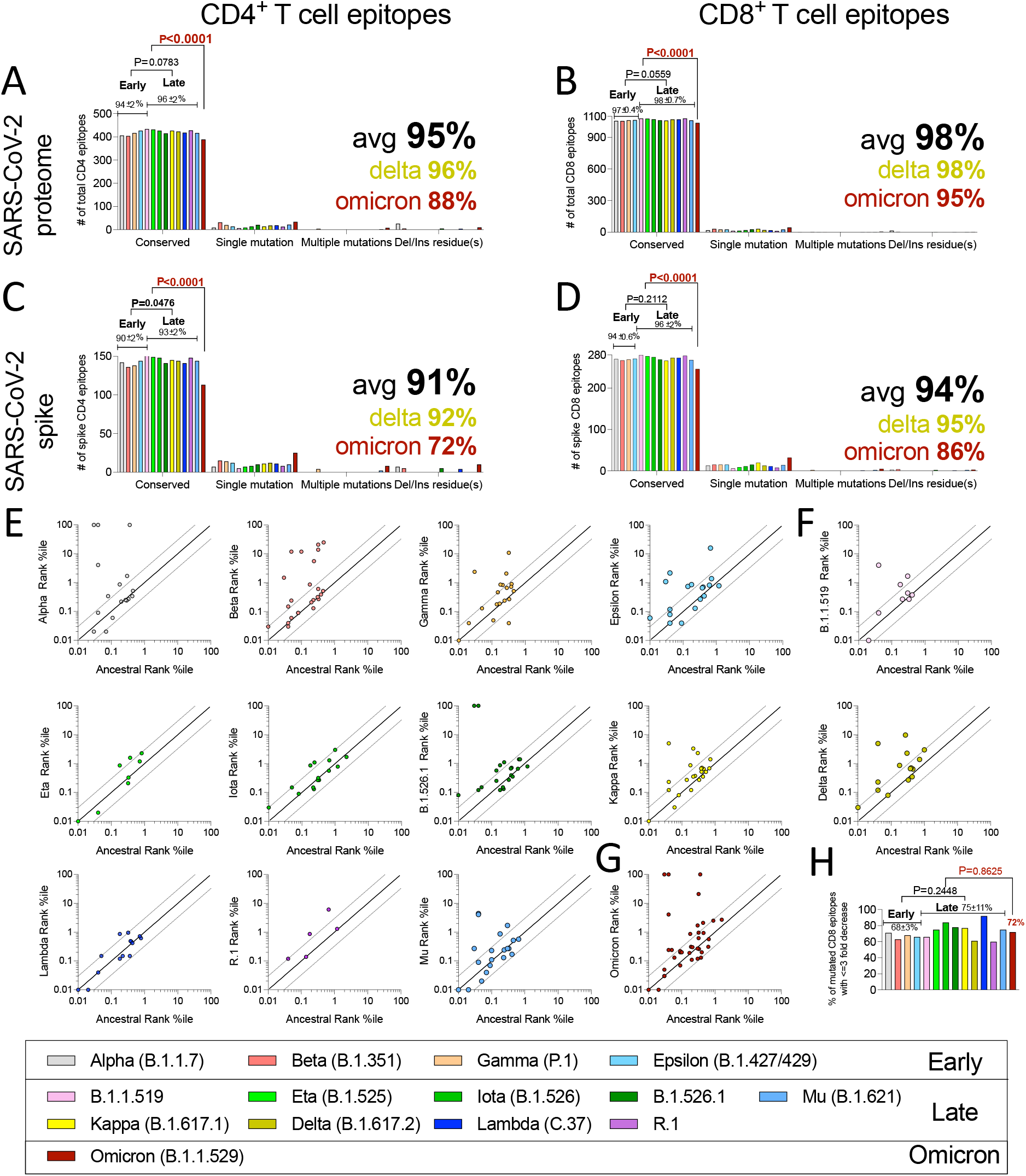
Conservation of ancestral SARS-CoV-2 T cell epitopes in a panel of SARS-CoV-2 variants. The number of epitopes that are fully conserved, or that are associated with single or multiple mutations, insertions/deletions was computed across SARS-CoV-2 variants. The analysis shown represents the breakdown of conserved and mutated CD4^+^ (**A, C**) and CD8^+^ epitopes (**B, D**) for all SARS-CoV-2 proteins (**A-B**) and spike protein only (**C-D**). The percentage of conserved epitopes was calculated for each variant separately. Average conservancy and standard deviations were calculated for all the variants, or separately for early variants, more recent SARS-CoV-2 variants, and Omicron. (**E-H**) The predicted HLA binding affinities of mutated versus ancestral sequences of CD8^+^ epitopes are shown, based on the epitope/HLA combinations curated in the IEDB data as of July 2021. Predicted HLA binding values to the relevant HLA allelic variant were calculated using the IEDB recommended NetMHCpanEL 4.1(Reynisson et al., 2020) algorithm. Points outside the dotted lines in each panel indicate instances where the predicted HLA binding capacity of the mutated peptide was increased (>3-fold) or decreased (<3-fold). (**E**) Early, (**F**) Late, and (**G**) Omicron SARS-CoV-2 variants are shown. (**H**) Percentage of mutated CD8^+^ T cell epitopes associated with a decrease of less than 3-fold in predicted binding capacity. Comparisons of epitopes conservancy across early and current variants are performed by unpaired Mann-Whitney test. Comparison with the B.1.1529 variant was performed by One sample T test. Large font bold numbers indicate average conservation in all variants (black), Delta (ochre) and Omicron (dark red). See also Table S3 and S4.

These values were conservative estimates of the number of preserved epitopes, since conservative substitutions and changes not impacting HLA binding can still be cross-reactively recognized. Accordingly, we examined the effect of the mutations on the predicted binding affinity of each CD8 epitope for which HLA restriction could be inferred (**Figure 4E-G**). Notably, in the majority of cases, the variant-associated mutations were predicted to not impair HLA binding capacity (**Figure 4E-G**). Importantly, 72% of the epitopes with Omicron variant mutations were predicted to retain similar HLA class I binding capabilities, which is not dissimilar to other SARS-CoV-2 variants (p=0.8625; **Figure 4H**). In conclusion, bioinformatic analyses suggest that the majority of CD4^+^ and CD8^+^ T cell epitopes are unaffected by mutations, regardless of whether early or late variants were considered (**Figure 4A-D**), thus suggesting that variant evolution was not driven by T cell escape. In the case of Omicron, the number of totally conserved epitopes is decreased, as expected on the basis of the larger number of spike mutations associated with Omicron. However, the majority of Omicron epitopes (full proteome or spike) were still 100% conserved, and the majority of mutated epitopes were predicted to still be recognizable by T cells.

### Experimental assessment of Omicron variants T cell mutations

Next, we experimentally determined the impact of Omicron mutations on T cell responses, in comparison to other variants, in a cohort of individuals vaccinated 5-6 months before donating blood and also in parallel subsequently used for epitope mapping. The overall conservation of memory CD4^+^ T cell recognition of Omicron spike was 0.84 by AIM and 0.75 by ICS assay (fold change). A significant decrease was observed for Omicron and the magnitude of the reduction was comparable to that of Alpha or Beta variants by AIM, while a significant decrease we observed only for Alpha by ICS (**Figure 5A-B**). At the individual subject level, no AIM^+^ CD4^+^ decreases >3-fold were observed. Cytokine-producing CD4^+^T cells decreases >3-fold were observed in 9 of 170 instances (5%) and no instances greater than 10-fold (**Figure 5B**). The conservation of memory CD8^+^ T cell recognition of Omicron spike was 0.85 by AIM (fold change) and 1.5 by ICS; both were not significant, while significant decreases were observed for Alpha, Beta and Delta by AIM (**Figure 5C-D**).

**Figure 5.**
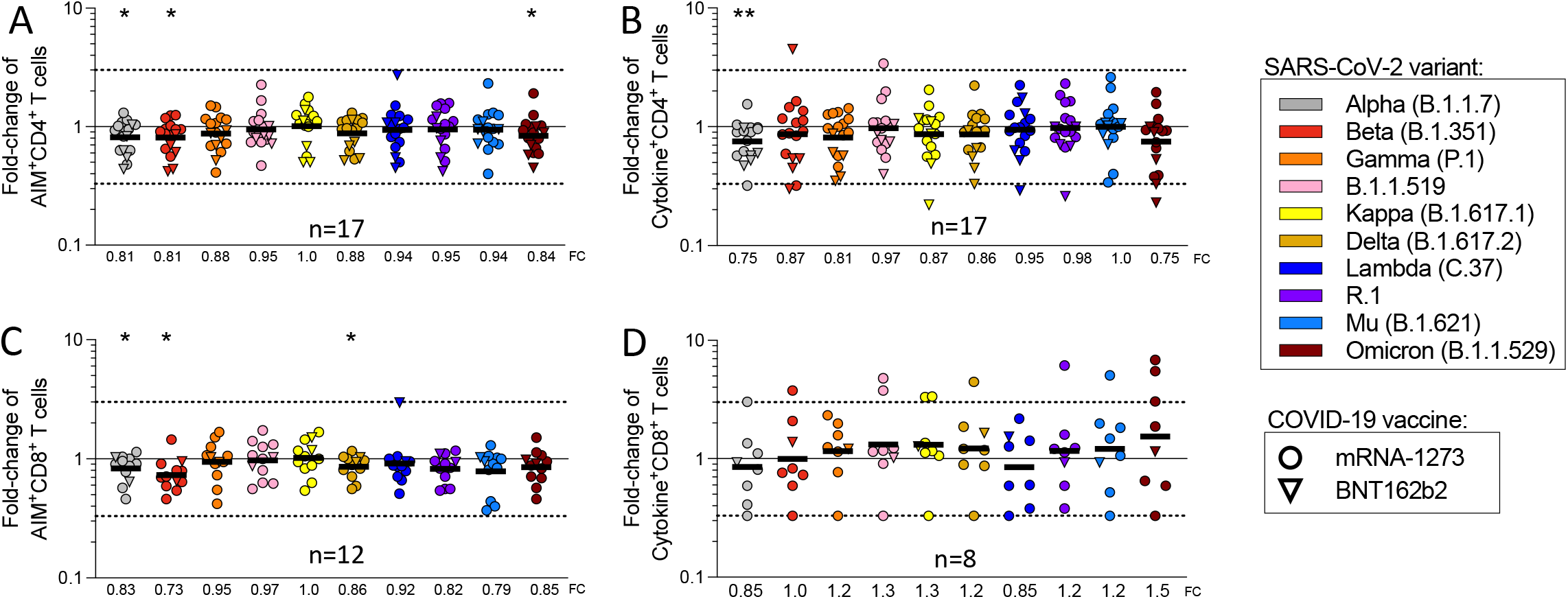
Impact of Omicron and other SARS-CoV-2 variants on memory T cell and B cell recognition. The response to SARS-CoV-2 variants was assessed in individuals 5-6 months after full vaccination with mRNA-1273 (circles) and BNT162b2 (triangles). The fold change values are shown for AIM^+^ (**A**) CD4^+^ (**B**) and CD8^+^ (**C**) T cell responses. (**C**) The Fold change of all cytokine^+^CD4^+^ T cells including IFNγ, TNFα, IL-2, Granzyme B, and CD40L. (**D**) The fold change of all cytokine^+^CD8^+^ T cells as calculated from the IFNγ^+^, TNFα^+^, IL-2^+^, or Granzyme B^+^ CD8^+^ T cells. Significance of fold change decreases for each variant was assessed by Wilcoxon Signed Rank T test compared to a hypothetical median of 1. See also Figure S5 and Table S1

For all timepoints analyzed in this study, we found a very weak inverse correlation between fold change decrease and the magnitude of the spike-specific T cell responses (**Figure S5E-F**), suggesting that overall weaker responses tend to be less frequently associated with decreases in the variants. This might simply reflect weaker responses being associated with a lesser dynamic range and therefore decreases less reliably measured. In any case, it did not support the notion that significant decreases are selectively associated with weak responses. We also examined the notion that weaker responses might be associated with individual HLA allele combinations, utilizing bioinformatic tools specifically designed to detect HLA associations (Paul et al., 2017). No specific HLA class I or class II alleles were significantly correlated to reduced variant recognition in our cohort (data not shown), but the limited sample size was not powered to detect HLA associations, which usually required substantially larger numbers of observations.

To further examine the molecular mechanism involved in the observed effects of T cell recognition of variant spike epitopes, we selected four donors for in depth spike epitope identification studies and variant analyses (**Figure 6A-D, Table S5**). Each vaccinated donor recognized 5 to 42 (median 11) individual CD4^+^ T cell epitopes in spike (**Figure 5A**). Approximately 80% of the CD4^+^ T cell response was associated with epitopes fully conserved in Omicron, with the actual values per donor ranging from 65% to 100% (**Figure 6B**). Each vaccinated donor recognized 6 to 19 (median 10) spike CD8^+^ T cell epitopes (**Figure 6C**). Approximately 80% of the CD8^+^ T cell response was associated with epitopes fully conserved in Omicron, with the values per donor ranging from 70% to 100% (**Figure 6D**). These results were in agreement with the bioinformatic analyses (**Figure 4**). In sum, these epitope mapping data show how the wide epitope repertoire associated with vaccine-induced responses counterbalances the effect of variant mutations of certain spike epitopes.

**Figure 6.**
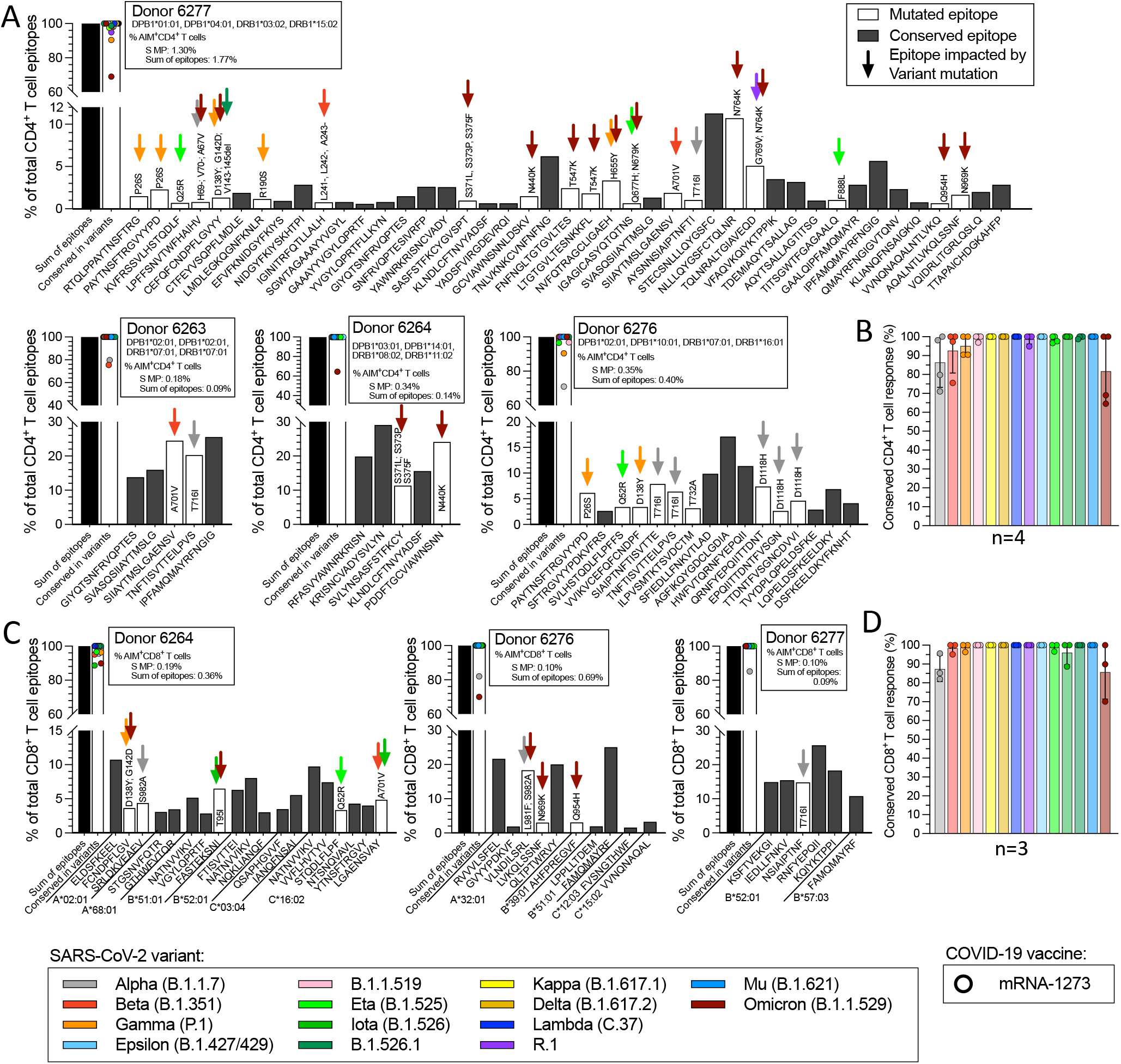
Impact of SARS-CoV-2 variants on spike epitope repertoires of fully vaccinated donors 5-6 months after vaccination. The response to SARS-CoV-2 variants was assessed in individuals 5-6 months after full vaccination with mRNA-1273 (circles), BNT162b2 (triangles) and Ad26.COV2.S (squares). (**A**) CD4^+^ and (**C**) CD8^+^ T cell epitope repertoires were determined for 4 mRNA-1273 vaccinees (no CD8 epitopes were identified for donor 6263). The percent of T cell response associated with conserved epitopes for each individual donor for (**B**) CD4^+^ and (**D**) CD8^+^ T cells is shown for each variant assessed. Each graph shows the total response detected with the ancestral spike MP, and the summed total response detected against each of the individual epitopes identified. The histograms show the % of the total response accounted from each epitope where black bars indicate non-mutated epitopes, while mutated epitopes are represented by open bars, with color coding further indicating which variant mutations are associated with the epitope. Based on these data the fraction of the total response to each variant that can be accounted for by nonmutated epitopes can be calculated, as also shown in the graph. See also Table S1, S2, S3 and S5.

## DISCUSSION

Here we analyzed adaptive immunity in vaccinated individuals to a comprehensive panel of SARS-CoV-2 variants, including Delta and Omicron, for multiple vaccines. Our data demonstrate that the vast majority of T cell epitopes are fully conserved, not only in the “early” variants previously analyzed (Collier et al., 2021; Geers et al., 2021; Keeton et al., 2021; Melo-Gonzalez et al., 2021; Riou et al., 2021; Tarke et al., 2021b), but also in newer variants, suggesting that the continued evolution of variants has not been associated with increased escape from T cell responses at the population level.

At the level of the full proteome, which is relevant for natural infection, 95% of reported class II and 98% of class I epitopes were fully conserved by computational analysis. In the case of Omicron, the fraction of epitopes that were fully conserved dropped to 88% for class II and 95% for class I epitopes in the whole proteome. Focused only on spike, relevant in the context of vaccination, 91% of class II and 94% of class I epitopes were still fully conserved. The fraction of totally conserved spike epitopes in Omicron dropped to 72% for class II and 86% for class I epitopes. The higher number of mutated T cell epitopes in spike was expected since many variant mutations are localized in the spike protein. Overall, the majority of T cell epitopes are conserved at the sequence level in all variants analyzed so far, including Omicron. It should be emphasized that an epitope mutation does not preclude cross-reactive recognition of the mutated sequence. To partially address this point, we calculated the fraction of class I epitope mutations predicted to be associated with a decrease in binding affinity to the relevant HLA. We found that, of the mutated epitopes, HLA binding was conserved for 72% of the epitopes. The impact on HLA binding was not different for Omicron epitopes compared to other variants. These observations argue against a model that mutations accumulated in Omicron might be the result of T cell immune pressure at the population level.

T cell recognition of several variants, including Delta and Omicron, was experimentally measured in donors vaccinated with mRNA-1273, BNT162b2 or Ad26.COV2.S. Variant recognition relative to the ancestral sequence was similar in the three different vaccine platforms tested, which is reassuring in terms of the potential implications for protective effects being similar regardless of the vaccine platform considered. A significant higher variability was detected with Ad26.COV2.S, but no clear explanation for this effect was apparent. A significant trend was also noted for more frequent decreases when utilizing the IFNγ production as a readout; while it is possible that this particular function is more impacted in mutated sequences, the measured outcomes may also be related to assay sensitivity limits.

A majority of memory T cells were not impacted by variants’ mutations, which is again reassuring in terms of the potential implications for T cell protective effects being similar regardless of the different vaccine cohorts considered. Memory T cell responses to the various variants, including Omicron, were dissected in detail in a cohort of donors 6-7 months following vaccination. The results confirmed that the majority of both CD4^+^ and CD8^+^ T cell responses detected by the AIM assay were preserved at this late time point. Some CD8^+^ T cell response decreases were observed when utilizing the IFNγ production as a readout. Of note, regardless of the assay, Omicron responses were largely preserved in both CD4^+^ and CD8^+^ T cells. In depth epitope identification experiments revealed that both CD4^+^ and CD8^+^ T cell responses in vaccinated donors were broad, and the data further demonstrated that for each individual donor/variant combination the majority of responses recognized epitopes 100% conserved. These data provide a clear explanation for the limited impact of variant associated mutations on T cell responses.

Adaptive immunity against SARS-CoV-2 consists of multiple branches (Sette and Crotty, 2021). Memory B cell recognition of variants’ spikes was reduced in all cases, but the reductions were < 3-fold, including against Delta spike and Beta RBD, demonstrating substantially retained memory B cell recognition of variants (Cho et al., 2021; Goel et al., 2021; Sokal et al., 2021a). This is consistent with the observations that neutralizing antibody titers against Omicron are generally low in individuals after two doses of mRNA-1273 or BNT162b2 but Omicron neutralizing antibody titers rapidly increase after a third immunization (Liu et al., 2021; Planas et al., 2021; Schmidt et al., 2021; Sokal et al., 2021b). Memory B cells may have important contributions in protective immunity by making anamnestic neutralizing antibody responses after infection (Cameroni et al., 2021; Carreño et al., 2021; Cele et al., 2021; Doria-Rose et al., 2021; Gagne et al., 2021; Garcia-Beltran et al., 2021; Zou et al., 2021).

This data provides reason for optimism, as most vaccine-elicited T cell responses remain capable of recognizing all known SARS-CoV-2 variants. Nevertheless, the data also underline the need for continued surveillance and the potential danger posed by continued variant evolution that could result in further reduction of T cell responses. Incorporation of additional elements eliciting broader T cell responses directed towards more conserved targets into vaccine strategies may be considered as a means to increase vaccine effectiveness against future variants.

## Limitations of the Study

The present study is associated with limitations, including the fact that memory B cells were not assessed for Omicron spike binding. A further caveat is that it is unknown what level of epitope conservation is likely to preserve functional T cell responses in vivo. Finally, the current study has not investigated subjects following natural infection.

## ACKNOWLEDGEMENTS

This project has been funded in whole or in part with Federal funds from the National Institute of Allergy and Infectious Diseases, National Institutes of Health, Department of Health and Human Services, under Contract No. 75N93021C00016 to A.S. and Contract No. 75N9301900065 to A.S, D.W. A.T. was supported by a PhD student fellowship through the Clinical and Experimental Immunology Course at the University of Genoa, Italy. We thank Gina Levi and the LJI clinical core for assistance in sample coordination and blood processing. We gratefully thank the authors from the originating laboratories responsible for obtaining the specimens, as well as the submitting laboratories where the genome data were generated and shared via GISAID, and on which this research is based. We would like to thank Vamseedhar Rayaprolu and Erica Ollmann Saphire for providing the recombinant SARS-CoV-2 Receptor Binding Domain (RBD) protein used in the ELISA assay.

## AUTHOR CONTRIBUTIONS

Conceptualization: A.G., S.C. and A.S.; Data curation and bioinformatic analysis, A.G., J.S..; Formal analysis: A.T., A.G., B.G, Z.Z, C.C., E.D.Y; Funding acquisition: S.C., A.S. D.W.; Investigation: A.T., N.M., Z.Z, C.H.C, N.I.B, E.D.Y, R.D.A and A.G.; Resources: S.M, E.P. Supervision: J.S., G.F.,D.W., S.C., A.S., R.A. J.D. and A.G.; Writing: A.G., S.C. and A.S.

## TABLES

## STAR METHODS

### RESOURCE AVAILABILITY

#### Lead Contact

Please refer to the Lead Contact (Alessandro Sette) for further information pertaining to availability of resources and reagents.

#### Materials Availability

Upon specific request and execution of a material transfer agreement (MTA) to the Lead Contact or to A.G., aliquots of the peptide pools utilized in this study will be made available. Limitations will be applied on the availability of peptide reagents due to cost, quantity, demand and availability.

#### Data and Code Availability

All the data generated in this study are available in the published article and summarized in the corresponding tables, figures and supplemental materials.

## EXPERIMENTAL MODEL AND SUBJECT DETAILS

### METHOD DETAILS

#### SARS-CoV-2 variants of concern selection and bioinformatic analysis

The genome sequences for the B.1.1.7, B.1.351, P.1. and B. 1.427/429 variants were selected as previously described (Tarke et al., 2021b). For the additional variants selected, the sequence variations in the variant viruses were derived by comparison with Wuhan-1 (NC_045512.2). All mutated amino acids in the different variants are outlined in **Table S3**. To determine the impact of the selected variants on T cell epitopes, CD4 and CD8 T cell epitopes were extracted from the IEDB database (www.IEDB.org)(Vita et al., 2019) on July 8^th^ 2021 using the following query: Organism: SARS-CoV2 (ID:2697049, SARS2), Include positive assays only, No B cells, No MHC assays, Host: Homo sapiens (human) and either MHC restriction type: Class I for CD8 epitopes or Class II for CD4 epitopes. Additional manual filtering was performed on the extracted datasets allowing only epitopes of 9-14 residues in size for class I and 13-25 residues for class II. This resulted in a total of 446 and 1092 epitopes for CD4 and CD8, respectively. The binding capacity of SARS-CoV-2 T cell epitopes, and their corresponding variant-derived peptides, was determined for their putative HLA class I restricting allele(s) in a smaller epitope subset (n=833) where information regarding allele restriction was available. Prediction analyses for class I were determined utilizing the NetMHCpan EL4.1 algorithm (Reynisson et al., 2020) implemented by the IEDB’s analysis resource(Dhanda et al., 2019; Vita et al., 2019). Predicted binding for class I analyses are expressed in terms of percentile. For each epitope-variant pair a fold change (FC) of affinities (variant /WT) was determined, corresponding values FC >3, indicating a 3-fold or greater decrease in affinity due to the mutation, were accordingly categorized as a decrease in binding capacity, and a FC <0.3 as an increase; FCs between 0.3 and 3 were designated as neutral.

#### Peptide synthesis and Megapool preparation

All the peptides used in this study were synthesized as crude material (TC Peptide Lab, San Diego, CA), and then individually resuspended in dimethyl sulfoxide (DMSO) at a concentration of 10– 20 mg/mL. For preparation of spike megapools sets of 15-mer peptides overlapping by 10 amino acids were synthetized to span the entire SARS-CoV-2 protein of the ancestral Wuhan sequence and a selection of the SARS-CoV-2 variants [AlphaB.1.1.7), Beta (B.1.351), Gamma (P.1), B.1.1.519, Kappa (B.1.617.1), Delta (B.1.617.2), Lambda (C37), R1, Mu (B.1.621) and Omicron (B.1.1.529)]. The Megapools (MP) for each variant were created by pooling aliquots of the corresponding individual peptides and then performing a sequential lyophilization. The resulting lyocake was subsequently resuspended in DMSO at 1 mg/mL as previously described (Grifoni et al., 2020; Tarke et al., 2021a; Tarke et al., 2021b).

#### Human Subjects, blood isolation and HLA typing

The La Jolla Institute for Immunology (LJI) Clinical Core recruited healthy adults who had received the first and, when applicable, second dose of a COVID-19 vaccination among the mRNA-1273, BNT162b2, Ad26.COV2.S or NVX-CoV2373 available vaccinations. At the time of enrollment in the study, all donors gave informed consent. The LJI Clinical Core facility has collected blood draws under IRB approved protocols (LJI; VD-214) when possible two weeks after each vaccine dose administered (timepoint 1 and timepoint 2) and/or 3.5 months after the last dose received (timepoint 3). All donors had their SARS-CoV-2 antibody titers measured by ELISA, as described below. Additional information on gender, ethnicity, age and timepoint of collection of the vaccinee cohorts are summarized in **Table S1**.

The LJI Clinical Core performed blood collection and sample processing based on SOPs previously established and described (Dan et al., 2021; Tarke et al., 2021a). Whole blood was collected in heparin coated blood bags, and the cellular fraction was separated from plasma by a centrifugation at 1850 rpm for 15 minutes. The plasma was consequently collected and stored at −20°C for serology assays, while the cellular fraction underwent density-gradient sedimentation to obtain the PBMCs using Ficoll-Paque (Lymphoprep, Nycomed Pharma, Oslo, Norway)(Grifoni et al., 2020). Isolated PBMCs were stored in liquid nitrogen in cryopreserved cell recovery media containing 90% heat-inactivated fetal bovine serum (FBS; Hyclone Laboratories, Logan UT) and 10% DMSO (Gibco) until cellular assays were performed. HLA typing was performed by an ASHI-accredited laboratory at Murdoch University (Western Australia) for Class I (HLA A; B; C) and Class II (DRB1, DRB3/4/5, DQA1/DQB1, DPB1), as previously described(Tarke et al., 2021a) (**Table S2**). Pheresis blood donations from an additional cohort of mRNA vaccinees were provided by the contact research organization (CRO) BioVIT and collected under the same IRB approval (VD-214) at LJI.

#### SARS-CoV-2 serology and PSV neutralization assay

SARS-CoV-2 serology was performed for all plasma samples collected as previously described(Rydyznski Moderbacher et al., 2020). Briefly, 1 ug/mL SARS-CoV-2 spike (S) Receptor Binding Domain (RBD) was used to coat 96-well half-area plates (ThermoFisher Cat#3690), which were then incubated at 4°C overnight. After blocking the plates the next day at room temperature for 2 hours with 3% milk in phosphate buffered saline (PBS) and 0.05% Tween-20, the heat-inactivated plasma was added for an additional 90-minute incubation at room temperature, followed by incubation with the conjugated secondary antibody. Plates were read on the Spectramax Plate Reader at 450 nm using the SoftMax Pro. For data analysis of SARS-CoV-2 serology, the limit of detection (LOD) was defined as 1:3 while the limit of sensitivity (LOS) was established based on uninfected subjects, using plasma from normal healthy donors that did not receive COVID-19 vaccination.

The SARS-CoV-2 pseudovirus (PSV) neutralization assay was performed for timepoint 3 samples as previously described(Mateus et al., 2021). Briefly, a monolayer of VERO cells (ATCC, Cat# CCL-81) was generated by seeding 2.5×10^4^ cells in flat clear-bottom black 96-well plates (Corning, Cat# 3904). Recombinant SARS-CoV-2-spike pseudotyped VSV-ΔG-GFP were generated with the specific amino acid mutations listed: D614G (WT), B.1.1.7 (Alpha; 69-70 deletion, 144 deletion, N501Y, A570D, D614G, P681H), B.1.351 (Beta; L18F, D80A, D215G, 241-243 deletion, K417N, E484K, N501Y, D614G, A71V), P.1 (Gamma; L18F, T20N, P26S, D138Y, R190S, K417T, E484K, N501Y, D614G, H655Y, T1027I, V1176F) and B.1.617.2 (Delta; T19R, F157-R158 deletion, L452R, T478K, D614G, P681R, D950N). Pretitrated recombinant virus for each variant were incubated with serially diluted human heat-inactivated plasma at 37°C for 1-1.5 hours. Confluent VERO cell monolayers were added and incubated for 16 hours at 37°C in 5% CO_2_ then fixed in 4% paraformaldehyde in PBS pH 7.4 (Santa Cruz, Cat# sc-281692) with 10 μg/ml Hoechst (Thermo Scientific, Cat#62249). Cells were imaged using a Cell Insight CX5 imager to quantify the total number of cells and infected GFP expressing cells to determine the percentage of infection. Neutralization titers (inhibition dose 50-ID50) were calculated using the One-Site Fit Log IC50 model in Prism 8.0 (GraphPad), and calibrated to WHO international standard (20/268). Samples that did not reach 50% inhibition at the lowest serum dilution of 1:20 were considered as non-neutralizing and were calibrated as 10.73 IU/mL.

#### Flow cytometry-based T cell assays

Activation Induced Marker (AIM) and Intra Cellular Staining (ICS) assays have been separately described in detail previously (Grifoni et al., 2020; Mateus et al., 2021; Tarke et al., 2021b). In this study, we performed both assays separately at timepoint 3, while we combined them for timepoints 1 and 2. To assess the best protocol for AIM+ICS assay, we carried out the three assays in parallel in the same samples (**Figure S1**). The best assay configuration to retain AIM marker expression and simultaneously detect cytokines required the addition of CD137 antibody to culture as described in detail below. **Figure S1** shows also the comparison of this AIM+ICS protocol with the classical AIM or ICS assays; no significant differences are observed amongst protocols, suggesting that the combined assay can be used to simultaneously detect AIM^+^ cells and the cytokine profile.

In all assays, PBMCs were cultured in the presence of SARS-CoV-2-specific (ancestral or variant) MPs [1 μg/ml] in 96-well U-bottom plates at a concentration of 1×10^6^ PBMC per well. As a negative control, an equimolar amount of DMSO was used to stimulate the cells in triplicate wells phytohemagglutinin (PHA, Roche, 1μg/ml) stimulated cells were used as positive controls. After incubation for 24 hours at 37°C in 5% CO_2_ cells were either stained for AIM markers only or an additional incubation of 4 hours was carried out by adding Golgi-Plug containing brefeldin A, Golgi-Stop containing monensin (BD Biosciences, San Diego, CA) and in the case of the AIM+ICS assay combined CD137 APC antibody was additionally added in culture (2:100; Biolegend Cat# 309810). In all assays, cells were stained on their surface for 30 min at 4°C in the dark. For AIM assays, cells were then acquired directly, while for both ICS and AIM+ICS assays, cells were additionally fixed with 1% of paraformaldehyde (Sigma-Aldrich, St. Louis, MO) permeabilized and blocked for 15 minutes followed by intracellular staining for 30 min at room temperature.

All samples were acquired on a ZE5 5-laser cell analyzer (Bio-Rad laboratories) and analyzed with FlowJo software (Tree Star Inc.). The gates for AIM or cytokine positive cells were drawn relative to the negative and positive controls for each donor. A representative example of the gating strategy for AIM, ICS or AIM+ICS assays is depicted in **Figure S1**. Specifically, lymphocytes were gated, followed by single cells determination. T cells were gated for being positive to CD3 and negative for a Dump channel including in the same colors CD14, CD19 and Live/Dead staining. The CD3^+^CD4^+^ and CD3^+^CD8^+^ were further gated based on OX40^+^CD137^+^ and CD69^+^CD137^+^ AIM markers, respectively. For ICS, CD3^+^CD4^+^ and CD3^+^CD8^+^ cells were further gated based on a combination of each cytokine (IFNγ, TNFα, IL-2, Granzyme B) with CD40L or FSC-A, respectively (**Figure S1**). To the total cytokine response and T cell functionality was calculated from Boolean gating of single cytokines or Granzyme B that was applied to CD3^+^CD4^+^ or CD3^+^CD8 cells. In the resulting data generated from the AIM and ICS T cell assays, the background was removed from the data by subtracting the average of the % of AIM^+^ or Cytokine^+^ cells plated in triplicate wells stimulated with DMSO. The Stimulation Index (SI) was calculated by dividing the % of AIM^+^ cells after SARS-CoV-2 stimulation with the average % of AIM^+^ cells in the negative DMSO control. An SI greater than 2 and a LOS of 0.03% or 0.04 % AIM^+^ CD4^+^ or CD8^+^ cells, respectively, after background subtraction was considered to be a positive response based on the median twofold standard deviation of T cell reactivity in negative DMSO controls. For ICS, an SI greater than 2 and a LOS of 0.01% or 0.02% ICS^+^ CD4^+^ or CD8^+^ cells, respectively, after background subtraction was considered to be a positive response based on the median twofold standard deviation of T cell reactivity in negative DMSO controls for all timepoints, except timepoint 4 that had an LOS of 0.01% for CD8^+^ T cells.

#### Flow cytometry-based B cell assays

Detection of antigen-specific B cells by flow cytometry was performed using B cell probes consisting of SARS-CoV-2 viral proteins conjugated with fluorescent streptavidin, as previously described by our group (Dan et al., 2021). Spike and RBD recombinant proteins used in this study are described in the Key Resource Table. Two separate flow cytometry panels were used to identify spike or RBD variants. To enhance specificity, identification of both WT spike and WT RBD B cells was performed using two fluorochromes for each protein, prior to gating on variant B cells. For that, biotinylated WT SARS-CoV-2 spike was incubated with Streptavidin in either BV711 (BioLegend, Cat# 405241) or BV421 (BioLegend, Cat# 405225) at a 20:1 ratio (~6:1 molar ratio) for 1 hour at 4°C. In a separate panel, biotinylated WT RBD was also conjugated with streptavidin BV711 (BioLegend, Cat# 405241) or streptavidin PE-Cy7 (BioLegend, Cat# 405206) in a 2.2:1 ratio (~4:1 molar ratio). The streptavidin-fluorochrome conjugates used to tetramerize the SARS-CoV-2 variant proteins are listed as follows: Alpha (B.1.1.7) spike BUV737 (BD bioscience, Cat# 612775), Alpha RBD BV785 (Biolegend, Cat# 613013); Beta (B.1.351) RBD, BUV615 (BD bioscience, Cat# 613013),Gamma (P.1) spike, BV785 (Biolegend, Cat# 405249), Gamma RBD, BV737(BD biosciences, Cat# 612775), Delta (B.1.617.2) spike and RBD, Alexa Fluor 647 (Thermo Fischer Scientific, Cat# S21374). Streptavidin PE-Cy5.5 (Thermo Fisher Scientific, Cat# SA1018) was used as a decoy probe to minimize background by eliminating SARS-CoV-2 nonspecific streptavidin-binding B cells. Seven million PBMCs were placed in U-bottom 96 well plates and stained with a solution consisting in 5μ of biotin (Avidity, catalog no. Bir500A) to avoid cross reactivity among probes, 20 ng of decoy probe, 416 ng of spike and 20.1 ng of RBD per sample, diluted in Brilliant Buffer (BD Biosciences, Cat# 566349) and incubated for 1 hour at 4°C, protected from light. After washing with PBS, cells from both spike and RBD panels were incubated with surface antibodies diluted in Brilliant Buffer, for 30 at 4°C, protected from light. Viability staining was performed using Live/Dead Fixable Blue Stain Kit (Thermo Fisher, Cat# L34962) diluted 1:200 in PBS and incubation at 4°C for 30 minutes. Acquisition was performed on Cytek Aurora and analyses were made using Flow Jo v. 10.7.1 (BD Biosciences). The frequency of Variants-specific memory B cells was expressed as a percentage of WT spike or RBD memory B cells (Singlets, Lymphocytes, Live, CD3− CD14− CD16− CD56−CD19+ CD20+ CD38int/−, IgD− and/or CD27+ spike or RBD BV711+, spike or RBD BV421+). PBMCs from a known positive control (COVID-19 convalescent subject) and an unexposed subject were included to ensure consistent sensitivity and specificity of the assay. Streptavidin PE-Cy5.5 (Thermo Fisher Scientific, Cat# SA1018) was used as a decoy probe to minimize background by eliminating SARS-CoV-2 nonspecific streptavidin-binding B cells.

## QUANTIFICATION AND STATISTICAL ANALYSIS

Data and statistical analyses were performed in FlowJo 10 and GraphPad Prism 8.4, unless otherwise stated. Statistical details of the experiments are provided in the respective figure legends and in each methods section pertaining to the specific technique applied. Data plotted in logarithmic scales are expressed as geometric mean. In all assays, fold change (FC) was calculated as the ratio of the variant pool/ ancestral pool for samples with a positive ancestral pool response. Significance of fold change decreases for each variant was assessed by Wilcoxon Signed Rank T test compared to a hypothetical median of 1.

---

**Table S1.**
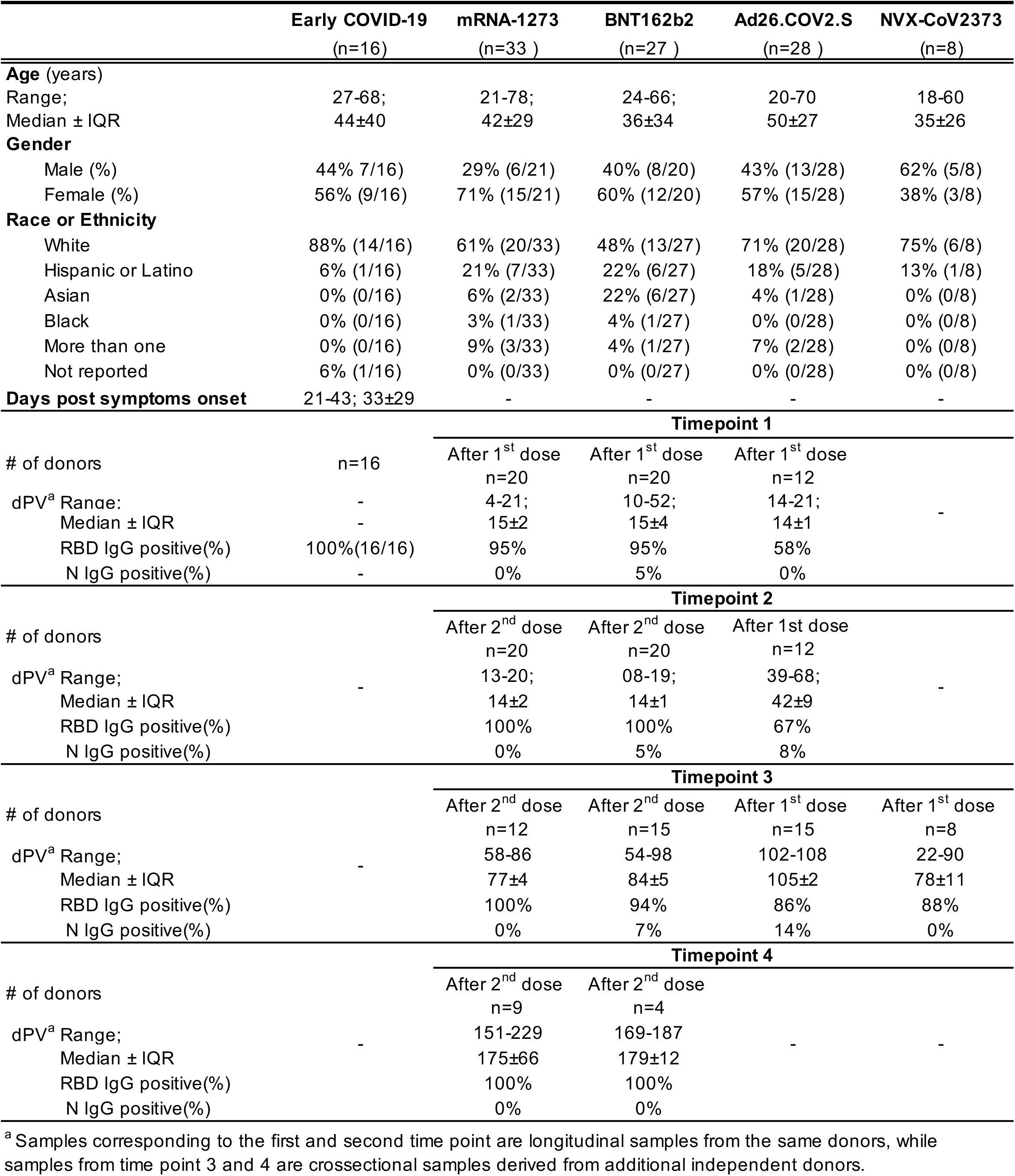
Related to Figures 1, 2, 3, 5 and 6. Characteristics of the donor cohorts enrolled in this study.

**Table S2.**
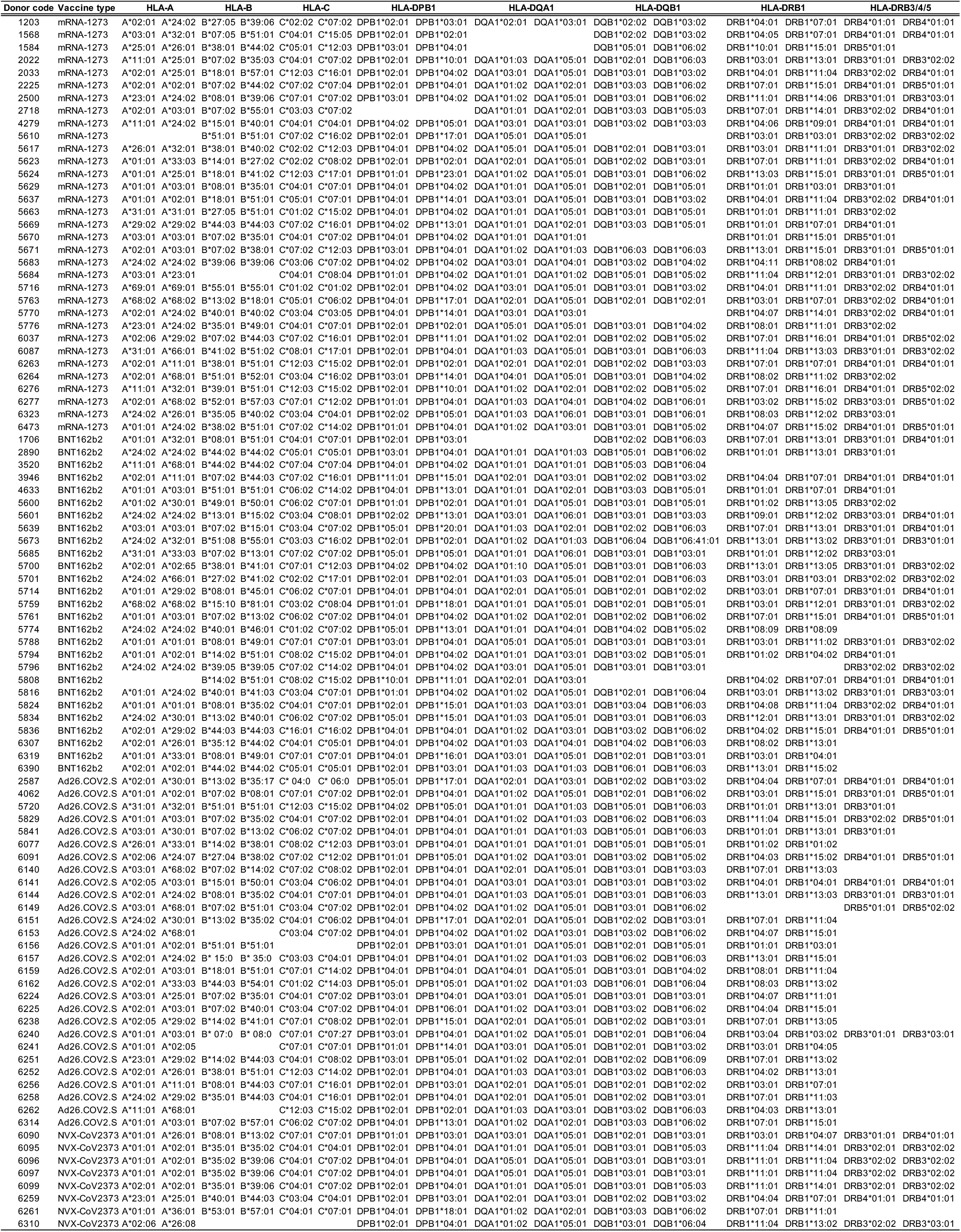
Related to Figures 1, 2, 3, 5 and 6. HLA typing of the cohort studied.

**Table S3.**
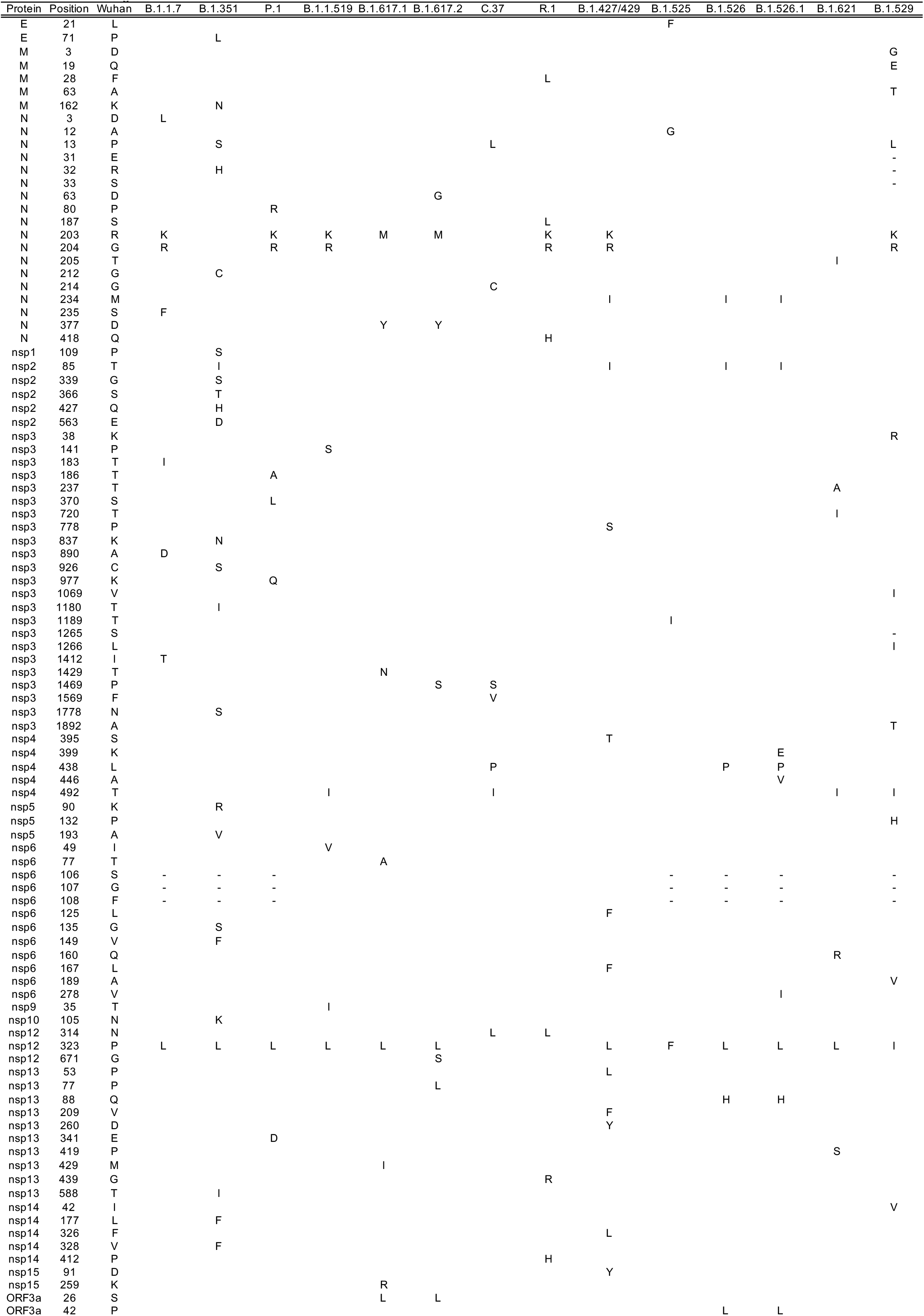
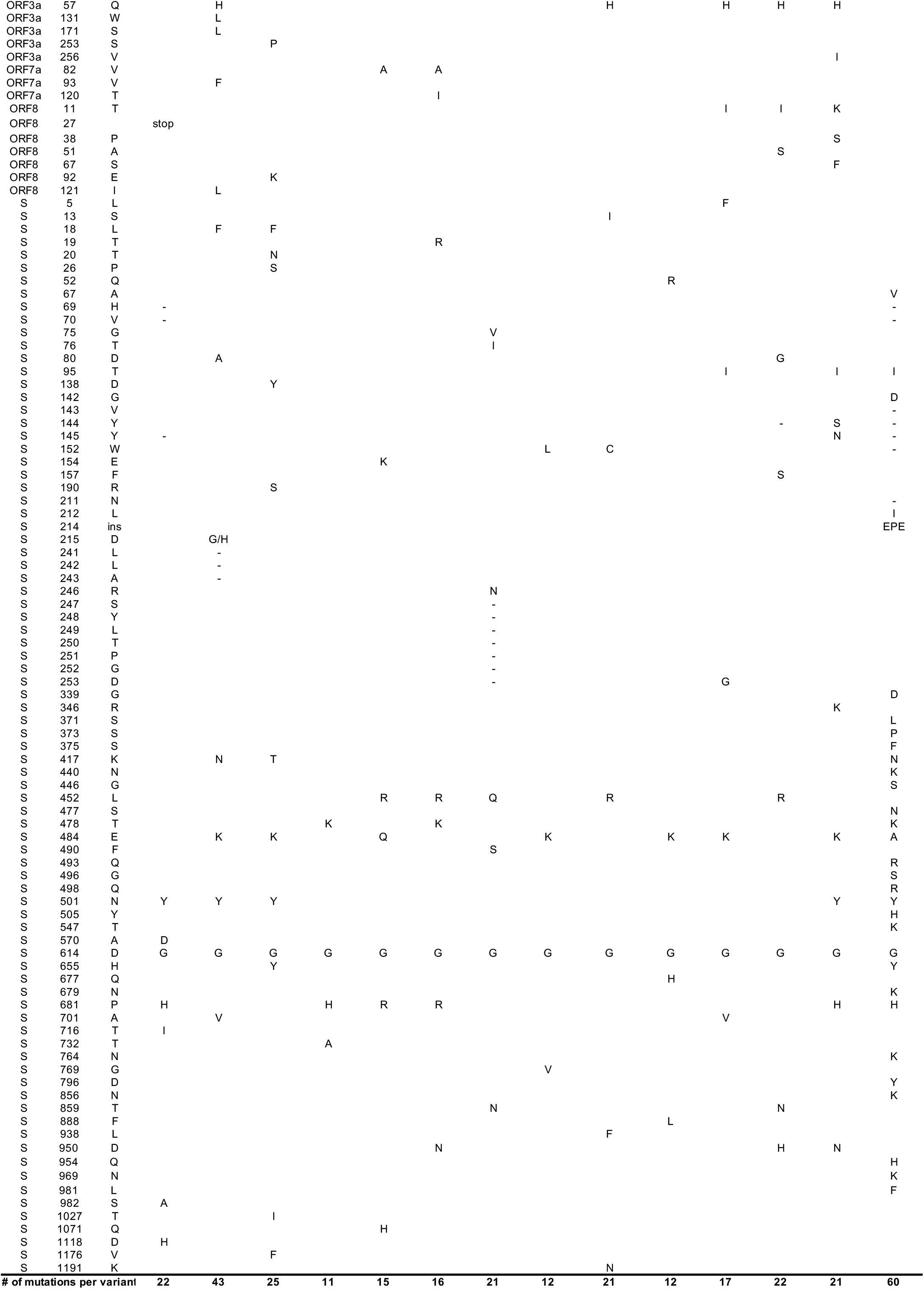
Related to Figures 1–6. List of amino acid mutations for each SARS-CoV-2 variant.

**Supplemental figure 1.**
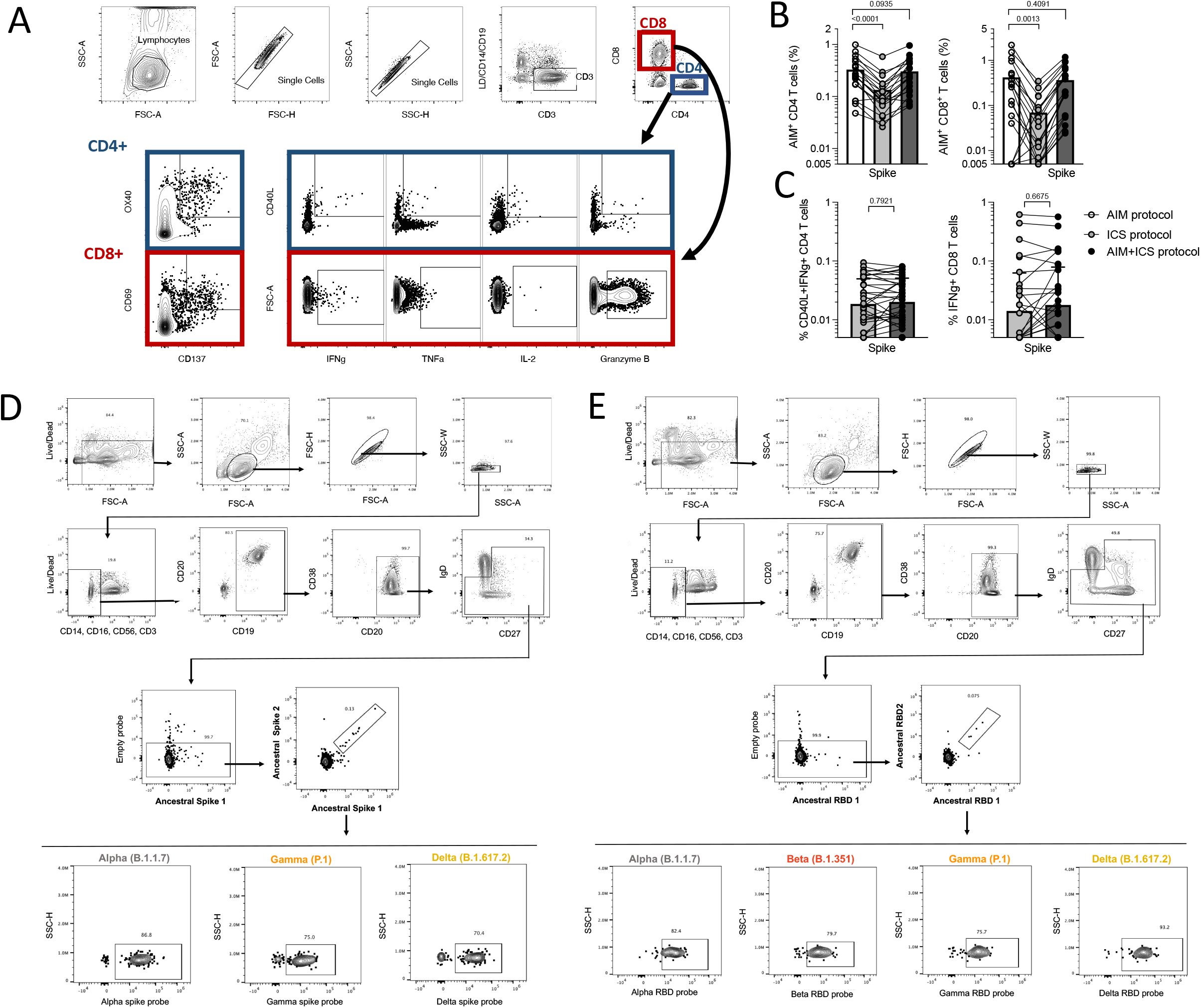
Related to Figures 1, 2, 3, 5 and 6. Assessment of SARS-CoV-2-specific T and B cells by flow cytometry based assays. (**A**) Gating strategy for T cell AIM, ICS, and AIM+ICS assays included in this study. Spike-specific responses are measured for both CD4^+^ and CD8^+^ T cells within the same donors using the indicated AIM markers or cytokines. (**B-C**) Validation of a combined AIM/ICS assay. The addition of a cocktail of Brefeldin and Monesin in the ICS assay significantly decreases the detection of AIM markers, while the inclusion of the CD137 antibody in culture concomitantly, repristinates the response (**B**) and does not impact the IFNγ detection (**C**). Data are shown after background subtraction and stimulation index > 2. Statistical analyses are performed using a paired Wilcoxon test. (**DE**) Representative gating strategy for the memory B cell assays using spike-protein or RBD, respectively.

**Supplemental Figure 2.**
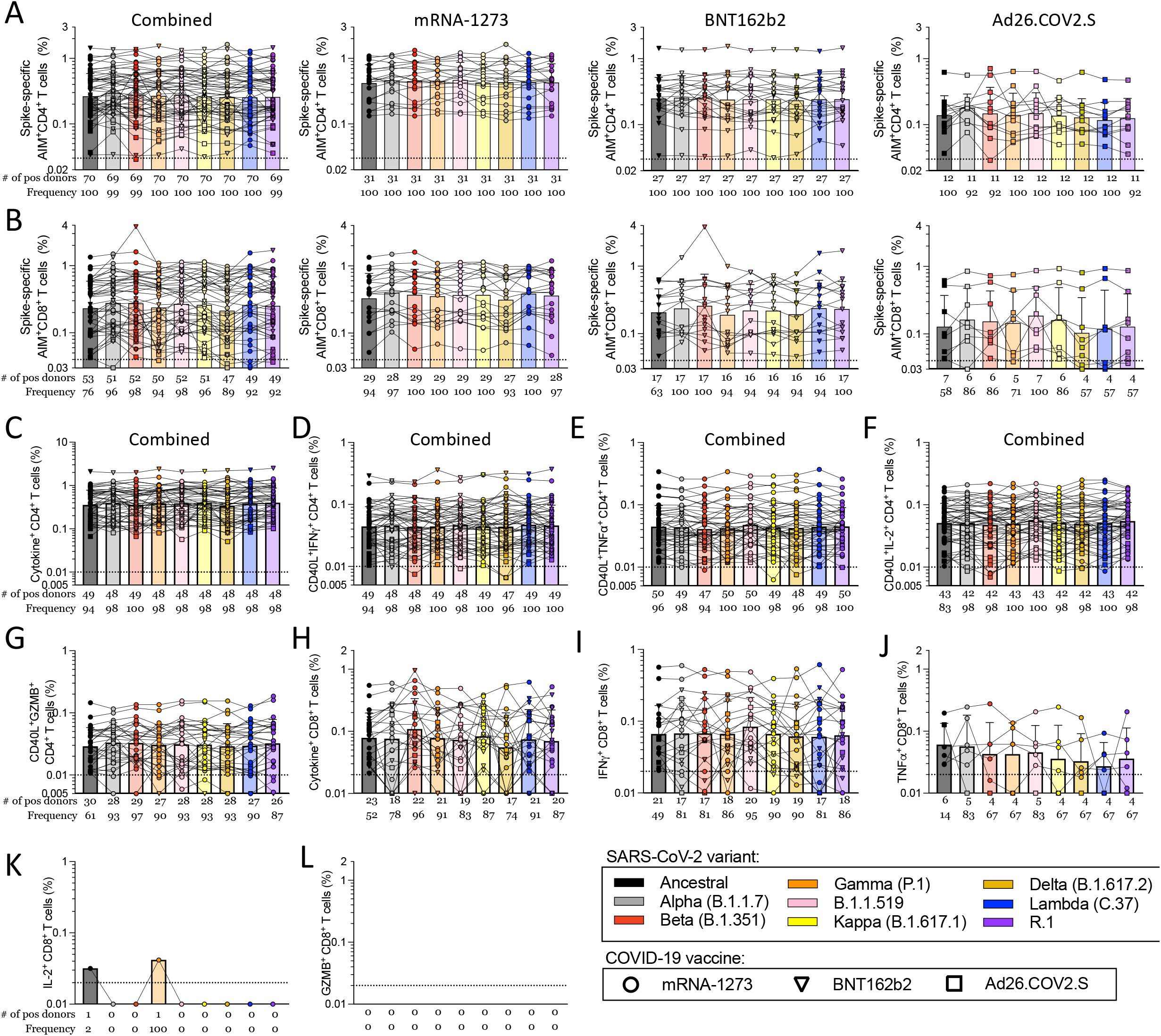
Related to Figures 1 and 2. Magnitude of CD4^+^ and CD8^+^ T cell responses in COVID-19 fully vaccinated individuals against ancestral and variant SARS-CoV-2 spike. AIM^+^ and cytokine^+^ T cell reactivities against MPs spanning the entire sequence of different SARS-CoV-2 variants are shown for PBMCs from fully vaccinated COVID-19 mRNA-1273 (circles), BNT162b2 (triangles) and Ad26.COV2.S (squares) vaccinees analyzed by vaccine platform or combined together. Data for (**A**) AIM^+^CD4^+^ and (**B**) AIM^+^CD8^+^ T cells is shown. The cytokine response of all vaccinees combined was quantified by (**C**) the total cytokine^+^CD4^+^ T cells which was calculated by measuring the cells expressing CD40L with (**D**) IFNγ, (**E**) TNFα, (**F**) IL-2, or (**G**) Granzyme B. For CD8^+^ T cells, the total cytokine response is shown (**H**) as calculated by the total IFNγ (**I**), TNFα (**J**), IL-2 (**K**), or Granzyme B (**L**) CD8^+^ T cells is shown. The frequency of response is based on the LOS (dotted line) for the ancestral response and SI>2, while the frequency of responses across different variants is based on the number of donors responding to the ancestral spike pool. All data shown is background subtracted.

**Supplemental Figure 3.**
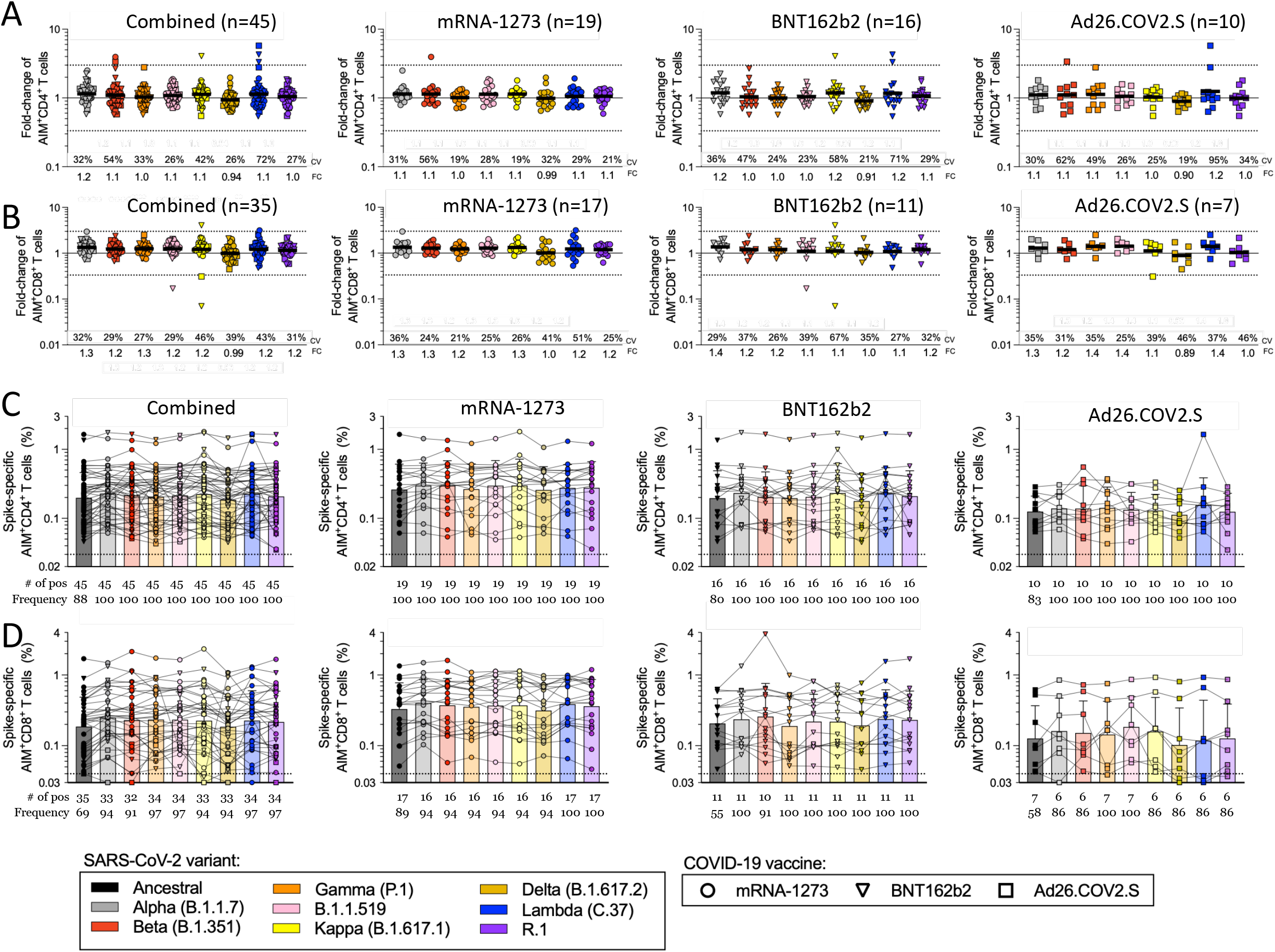
Related to Figure 1. Fold-change values and magnitude of AIM+ T cell responses 2 weeks after 1^st^ vaccination dose. CD4^+^ and CD8^+^ T cell responses of PBMCs were assessed with variant Spike MPs 2 weeks after the donors received the first dose of vaccination. The effect of mutations associated with each variant MP is expressed as relative (fold change variation) to the T cell reactivity detected with the ancestral strain MP. COVID-19 mRNA-1273 (circles), BNT162b2 (triangles) and Ad26.COV2.S (squares) vaccinees are presented combined together, and separately by vaccine platform. The fold-change is calculated in respect to the ancestral strain in COVID-19 vaccinees for (**A**) AIM^+^ CD4^+^ and (**B**) AIM^+^ CD8^+^ T cells. The magnitude of AIM^+^ T cell reactivity against the spike pools is shown for (**C**) CD4^+^ and (**D**) CD8^+^ T cells. The frequency of response is based on the LOS (dotted line) for the ancestral response and SI>2, while the frequency of responses across different variants is based on the number of donors responding to the ancestral spike pool. Coefficients of variation (CV) and geometric mean of the fold change (FC) for the variants are listed in each graph. Significance of fold change decreases for each variant was assessed by Wilcoxon Signed Rank T test compared to a hypothetical median of 1.

**Supplemental Figure 4.**
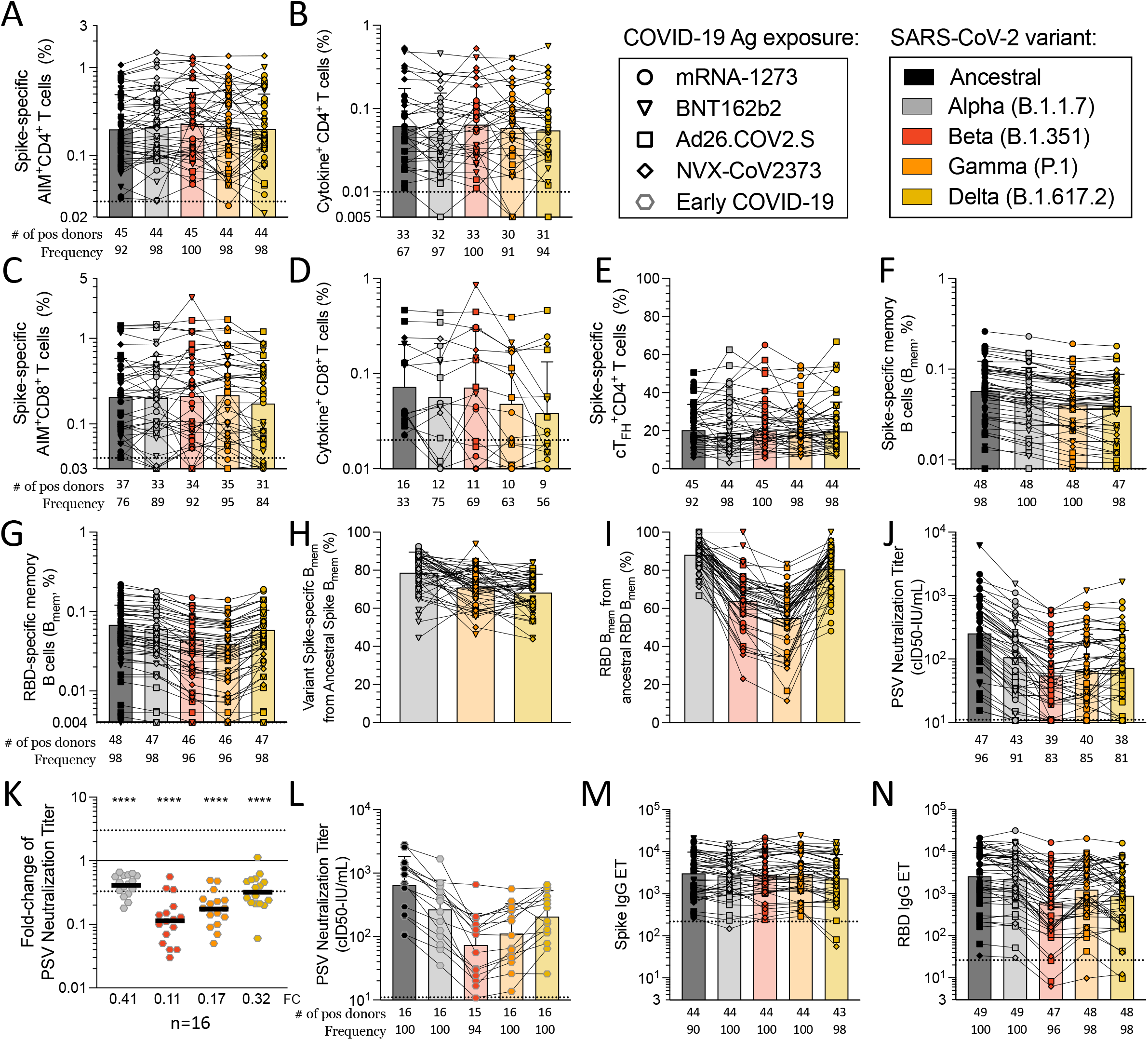
Related to Figure 3. Magnitude of T and B cell responses in COVID-19 vaccinated individuals 3.5 months after vaccination and antibody neutralization titer with early COVID-19 infected individuals. COVID-19 mRNA-1273 (circles), BNT162b2 (triangles), Ad26.COV2.S (squares) and NVX-CoV2373 (diamonds) vaccine recipients were assessed for T and B cell responses to variant SARS-CoV-2 spike MPs and all vaccine platforms are analyzed together. The magnitude of response is shown for (**A**) CD4^+^ T cells in the AIM assay and (**B**) the sum cytokine^+^CD4^+^ T cells, calculated from CD40L^+^, IFNγ^+^, TNFα^+^, IL-2^+^, or Granzyme B^+^ cells. The magnitude of responding CD8^+^ T cells are shown for (**C**) the AIM assay and (**D**) the sum of cytokines calculated from the CD8^+^ cells with intracellular expression of IFNγ, TNFα, IL-2, or Granzyme B. (**E**) The total magnitude of spike-specific AIM^+^ cT_FH_^+^CD4^+^ T cells is shown. The frequency of (**F**) Spike- and (**G**) RBD-specific B cells among total memory B (B_mem_) cells was assessed, as well as the frequency of variant-specific B_mem_ response within the ancestral response to (**H**) Spike and (**I**) RBD. (**J**) The antibody neutralization assay titer is shown for COVID-19 vaccinees. (**K**) The fold change values are shown for early COVID-19 infected donors for the neutralization assay and (**L**) the magnitude of the neutralization titers for these donors. (**M**) Spike and (**N**) RBD IgG titers are shown. The frequency of response is based on the LOS (dotted line) for the ancestral response and SI>2, while the frequency of responses across different variants is based on the number of donors responding to the ancestral spike pool. Significance of fold change decreases for each variant was assessed by Wilcoxon Signed Rank T test compared to a hypothetical median of 1.

**Supplemental Figure 5.**
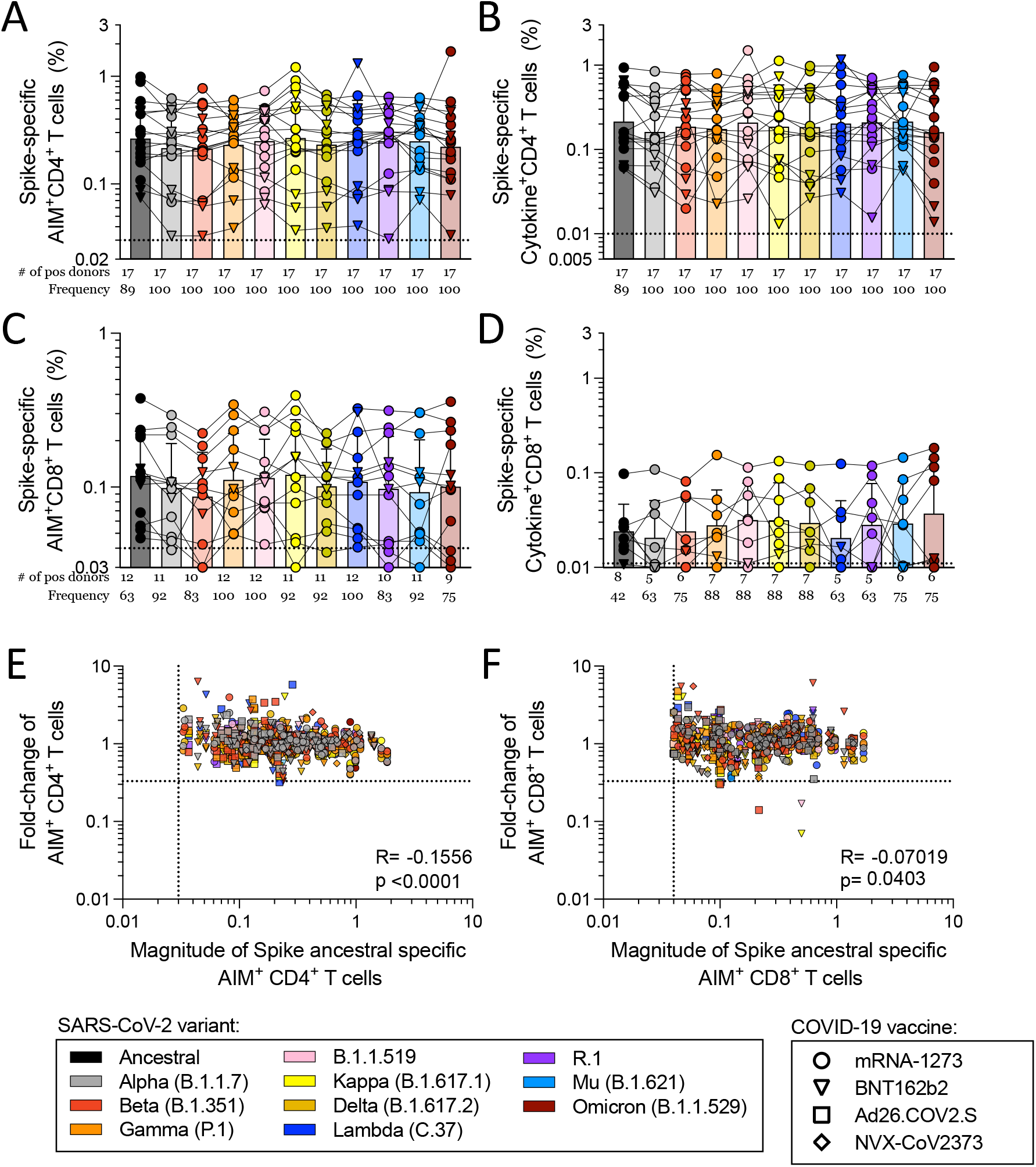
Related to Figure 5. Response to SARS-CoV-2 variants in fully vaccinated donors 5-6 months after vaccination. 5-6 months after vaccination, COVID-19 mRNA-1273 (circles) and BNT162b2 (triangles) vaccine recipients were assessed for T cell responses to variant Spikes by AIM and ICS assays. (**A**) The magnitude of response is shown for AIM+CD4+. (**B**) The sum of CD4^+^ CD40L^+^, IFNγ^+^, TNFα^+^, IL-2^+^, or Granzyme B+ T cells is shown. For CD8^+^ T cells, (**C**) the magnitude of AIM^+^CD8^+^ T cells is shown and (**D**) the total cytokine^+^CD8^+^ T cells calculated by summing the IFNγ^+^, TNFα^+^, IL-2^+^, or Granzyme B^+^ CD8^+^ T cells. COVID-19 mRNA-1273 (circles), BNT162b2 (triangles), Ad26.COV2.S (squares), and NVX-CoV2373 (diamonds) vaccine recipients were assessed for T cell responses to variant Spikes by AIM assay at various timepoints ranging from 2 weeks after the first dose to 5-6 months after the last dose of vaccination. The correlation of magnitude and fold-change values (**E**) AIM^+^ CD4^+^ or (**F**) CD8+ T cells was analyzed for all timepoints combined. R and p values are the results of a Pearson correlation.

**Table S5.**
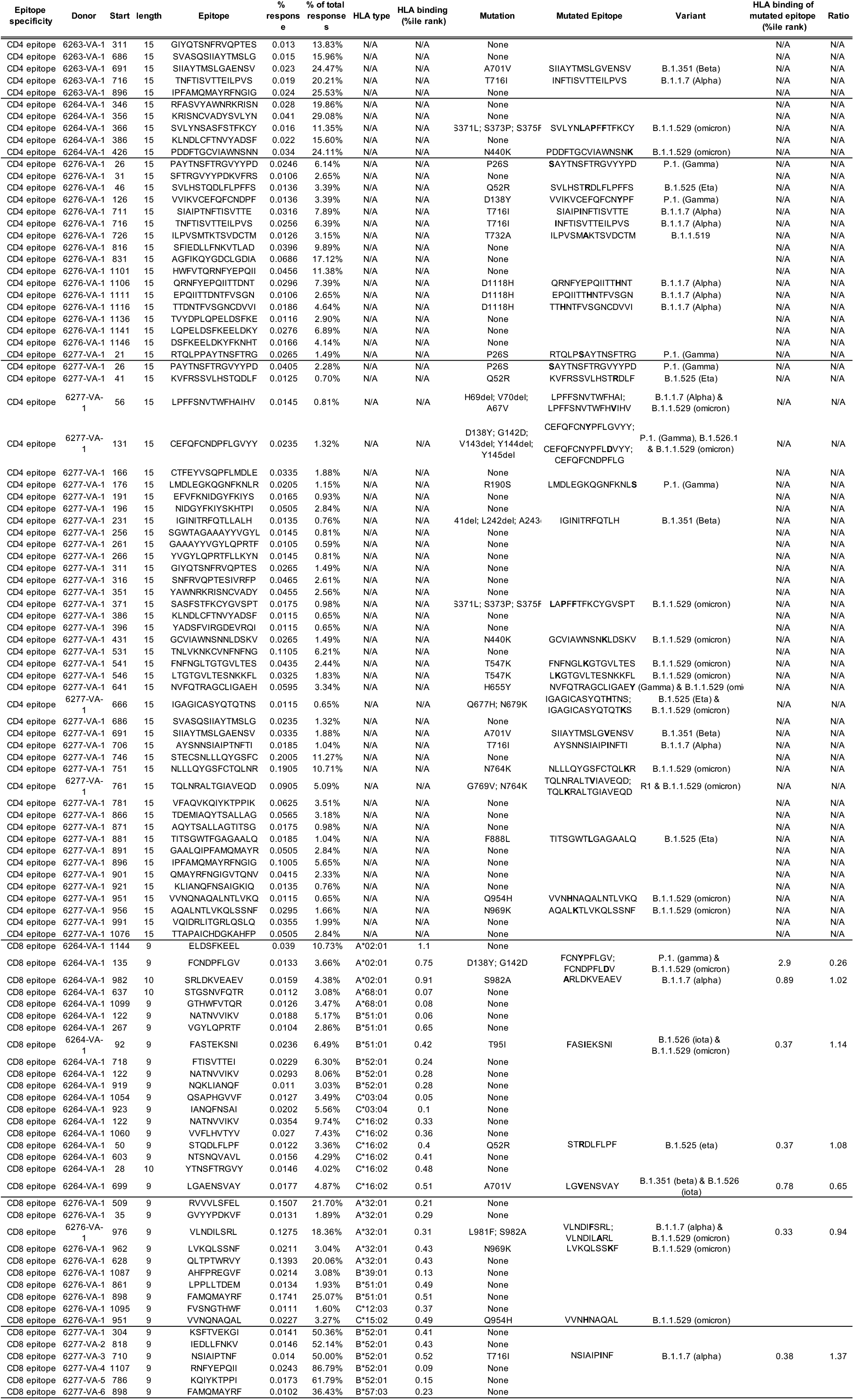
Related to Figure 6. List of class II and class I epitopes identified in vaccinated apheresis donors. Information on the mutations within these epitopes when applicable and to which variants they belong are included.

